# Maintenance of High Inbreeding Depression in Selfing Populations: Effects of Coupling of Early- and Late-Acting Mutations

**DOI:** 10.1101/748699

**Authors:** Satoki Sakai

**Affiliations:** Department of Ecology and Evolutionary Biology, Graduate School of Life Sciences, Tohoku University, Sendai 980-8578, Japan

**Keywords:** inbreeding depression, selfing, deleterious mutation, early-acting locus, late-acting locus

## Abstract

High estimates of inbreeding depression have been obtained in many plant populations with high selfing rates. However, deleterious mutations might be purged from such populations as a result of selfing. I developed a simulation model assuming the presence of mutations at two sets of loci, namely, early- and late-acting loci, and the selective abortion of embryos coupled with ovule overproduction. In the model, early-acting loci are expressed during embryo initiation, and less vigorous embryos are aborted. Late-acting loci are expressed after selective abortion ends; the surviving embryos (seeds) compete, and some of them form the next generation. If mutations are allowed to occur in both early- and late-acting loci, they increase in frequency in populations with high selfing rates in both sets of loci. However, this phenomenon does not occur if mutations occur in only the early- or late-acting loci. Consistent results are observed even if the total number of loci in which mutations are allowed to occur is the same among simulations with both early- and late-acting loci or only early- or late-acting loci, indicating that the presence of both sets of loci is the causal factor. Thus, the coupling effects of early- and late-acting mutations promote the maintenance of these mutations in populations with high selfing rates.

---

In organisms in which selfing is possible, such as hermaphroditic plants, the offspring produced by selfing are often less vigorous (as a result of inbreeding depression). Such inbreeding depression is caused by recessive or partially recessive deleterious mutations (CHARLESWORTH AND CHARLESWORTH 1999; CHARLESWORTH AND WILLIS 2009).

Selfing is expected to purge deleterious mutations from a population because embryos that are homozygous for deleterious mutations are produced, and they will disappear. However, substantial variation in inbreeding depression exists among plants with similar selfing rates, with estimates ranging from nearly 0 to approximately 0.9 in plants with selfing rates of 0 to 0.8 and from 0 to approximately 0.5 in plants with selfing rates greater than 0.8 (WINN *et al*. 2011). Furthermore, BALDWIN AND SCHOEN (2019) experimentally forced selfing in self-incompatible populations and found that inbreeding depression was difficult to purge by selfing. Thus, many plants with high selfing rates maintain deleterious mutations. This discrepancy is a current problem in evolutionary biology. Furthermore, as explained below, the maintenance of deleterious mutations is strongly related to two other topics in evolutionary biology.

One of these topics is the evolution from outcrossing to selfing. This transition is a very common transition in flowering plants and has independently occurred in many plant taxa (STEBBINS 1974; BARRETT 2002). Selfing has two major short-term advantages. First, plants can produce seeds by selfing under pollinator- and/or mate-limitation conditions (DARWIN 1876; GOODWILLIE *et al*. 2005). Second, an allele promoting selfing has a transmission advantage over an allele promoting outcrossing because the former allele can be transmitted via the pollen parent of seeds produced by selfing (FISHER 1941; GOODWILLIE *et al*. 2005; ECKERT *et al*. 2006). However, if many deleterious mutations are maintained in selfing populations, these short-term advantages may be insufficient to outweigh the inbreeding depression in seeds produced by selfing. Furthermore, genetic load is a severe long-term disadvantage of selfing and can lead to the extinction of selfing populations (GOLDBERG *et al*. 2010; CHEPTOU 2019). An outcrossing population that maintains many deleterious mutations and begins to self may become extinct as a result of inbreeding depression. However, extinction may be avoided if deleterious mutations are rapidly purged. Therefore, understanding whether deleterious mutations are maintained in selfing populations is important.

The second topic is the evolution of selfing rates, which has been studied theoretically and empirically (LLOYD 1979; LANDE AND SCHEMSKE 1985; SCHEMSKE AND LANDE 1985; HOLSINGER 1991; UYENOYAMA AND WALLER 1991; LLOYD 1992; LATTA AND RITLAND 1994; CHEPTOU AND MATHIAS 2001; GOODWILLIE *et al*. 2005; PORCHER AND LANDE 2005a; PORCHER AND LANDE 2005b; WINN *et al*. 2011; LANDE AND PORCHER 2015). In particular, the conditions that select for intermediate selfing rates have been extensively studied. Although intermediate selfing rates are commonly observed in plants (GOODWILLIE *et al*. 2005; WINN *et al*. 2011), general theory predicts that populations will evolve to complete outcrossing or selfing dependent on the degree of inbreeding depression (LLOYD 1979; LANDE AND SCHEMSKE 1985). In addition, if selfing purges deleterious mutations, then complete selfing should be evolutionarily stable (LANDE AND SCHEMSKE 1985). Hence, whether deleterious mutations are maintained or purged affects the evolution of selfing rate.

The selective interference hypothesis is a convincing hypothesis that has been proposed to explain the maintenance of deleterious mutations (GANDERS 1972; LANDE *et al*. 1994). This hypothesis states that if nearly all embryos produced by selfing are homozygous for highly deleterious mutations at certain loci and die without developing to the next generation, then no opportunity exists to select for heterozygous embryos or embryos without mutations. Although there is some evidence consistent with this hypothesis, whether selective interference alone can explain the maintenance of deleterious mutations is unclear (WINN *et al*. 2011). Additionally, for this mechanism to operate, most mutations must be recessive; hence, selective interference might be unlikely to maintain inbreeding depression (KELLY 2007; ROZE 2015).

Two previous studies examined the maintenance of deleterious mutations assuming early- and late-acting genes. The effect of the overproduction of ovules on the maintenance of deleterious mutations is analyzed in Porcher and Lande (2005b). They assumed that the same mutations become both early acting and late acting; the effects appear during embryo development (i.e., are early acting) if the mutations are homozygous but appear later (i.e., are late acting) if the mutations are heterozygous. Thus, early-acting and late-acting genes are assumed not to differ. They showed that the overproduction of ovules (i.e., the production of more ovules than of seeds by the parent) promoted the maintenance of deleterious mutations. Lande and Porcher (2017) analyzed the effects of the interactions between early- and late-acting loci on the maintenance of deleterious mutations. For late-acting loci, they assumed quantitative genetic variance under stabilizing selection without considering the individual late-acting loci. They showed that there is a purging threshold rate of self-fertilization, before which the population is effectively outcrossing and deleterious mutations are maintained.

In the present paper, I propose a new mechanism by which deleterious mutations are maintained without being purged from populations with high selfing rates: the coupling effects of early-acting and late-acting loci. I also develop a model considering early-acting and late-acting loci. However, in my model, early- and late-acting loci are distinguished, and the effects of early- and late-acting mutations appear during their respective stages. (Whether they are homozygous or heterozygous does not affect the stage at which their effects appear.) In other words, early-acting loci are expressed during the initiation of embryos to the stage at which selective abortion of embryos ends (assuming overproduction of ovules), whereas late-acting loci are expressed only after selective abortion ends; these loci act during seed completion, seed germination, and seedling growth. Then, I consider inbreeding depression caused by early-acting and late-acting loci. Ovules are overproduced and therefore do not all develop into seeds mainly due to resource limitation, which is a very common phenomenon in plants (LEE 1988). I assume individual loci for both early- and late-acting loci and examine the effects of carrying both early- and late-acting mutations by the same embryos (coupling effects of mutations).

## Model

### Simulation process

I carried out simulations to examine when deleterious mutations were maintained in populations that underwent mutations at early- and late-acting loci. Each population consists of *n* diploid hermaphroditic plants with a selfing rate *s* and random outcrossing among *n* plants according to a prior selfing model (LLOYD AND SCHOEN 1992), as explained below. There is no pollen limitation, and all ovules produced are successfully fertilized by self or nonself pollen. I assumed nonoverlapping generations, with the parents in the new generation selected from the seeds produced by the parents in the previous generation. After the initial generation, mutations occur at rate *m*, which represents the number of mutations per gene per generation. Inbreeding depression may occur if deleterious mutations become homozygous through inbreeding. The simulations followed the dynamics of the deleterious mutations for 500 generations.

The symbols used in the model are defined in Table 1.

**Table 1.** Symbols used in the present model and their definitions.

| Symbol | Definition |
| --- | --- |
| $d_e$ | Selection coefficient against an individual mutation when homozygous for early-acting mutations |
| $d_l$ | Selection coefficient against an individual mutation when homozygous for late-acting mutations |
| $d_t$ | Threshold for the fitness values of embryos above which embryos can potentially develop to seeds |
| $h_e$ | Dominance coefficient for the early-acting loci |
| $h_l$ | Dominance coefficient for the late-acting loci |
| $hm_e$ | Numbers of homozygotes in the early-acting mutations |
| $ht_e$ | Numbers of heterozygotes in the early-acting mutations. |
| $hm_l$ | Numbers of homozygotes in the late-acting mutations |
| $ht_l$ | Numbers of heterozygotes in the late-acting mutations |
| $i$ | Number of chromosomes carrying both early- and late-acting loci in each haplotype genome (carried by the first to $i$th chromosomes) |
| $j$ | Number of early-acting loci in the chromosomes carrying those loci (with the first to $j$th loci in each chromosome being the early-acting ones) |
| $m$ | Mutation rate per gene per generation |
| $n$ | Number of plants in the population |
| $n_{\text{link}}$ | Number of chromosomes in each haplotype genome |
| $n_{\text{locus}}$ | Number of loci in each chromosome |
| $n_p$ | Number of pollen grains produced by a parent |
| $n_{\text{pm}}$ | Number of pollen mother cells ($n_p = 4n_{\text{pm}}$) |
| $n_o$ | Number of ovules produced by a parent |
| $n_s$ | Number of seeds a parent can potentially produce |
| $s$ | Selfing rate |

#### Early- and late-acting loci

Early- and late-acting loci are defined as described in the introduction.

#### Genomes

Each haplotype genome consists of *n*_link_ chromosomes, each with the same number of loci (*n*_locus_). Of the *n*_link_ chromosomes, the first to the *i*th chromosomes carry both early- and late-acting loci, where the first to the *j*th loci in each chromosome are early-acting loci, and the others are late-acting loci. The other chromosomes carry only late-acting loci.

#### Pollen and ovule production

Each parent produces *n*_p_ pollen grains (*n*_p_ = 4*n*_pm_, where *n*_pm_ is the number of pollen mother cells) and *n*_o_ ovules. Deleterious mutations occur during production of the megaspore and pollen mother cells, and the number of mutations in each mother cell is sampled from a Poisson distribution with the parameter 4*mn*_link_*n*_locus_. Recombination may occur during meiosis at rate *r* per chromosome per generation. If recombination occurs, a crossover between one pair of chromatids takes place at a randomly determined point (loci in the model are equidistant in cM). Furthermore, different chromosomes randomly segregate during meiosis. As a result, a pollen mother cell develops into 4 pollen grains, whereas one randomly selected megaspore, which develops from a megaspore mother cell, remains in the ovule.

#### Selfing and outcrossing

Within each parent, randomly selected *sn*_o_ ovules are fertilized by randomly selected *sn*_o_ self-pollen grains before outcrossing. The remaining *n*_p_ – *sn*_o_ pollen grains contribute to the population’s pollen pool. Outcrossing then occurs among all the *n* parents in the population. There is no pollen limitation, and the (1 – *s*)*n*_o_ ovules remaining within each parent after selfing are fertilized by pollen that is randomly selected from the pollen pool.

#### Early selection

Deleterious effects are assumed to be the same for all mutations at any early-acting locus. The relative fitness values are 1 – *d*_e_ (0 ≤ *d*_e_ ≤ 1) for homozygotes and 1 – *h*_e_*d*_e_ (0 ≤ *h*_e_ ≤ 1) for heterozygotes, where *d*_e_ is the selection coefficient against an individual mutation when homozygous, and *h*_e_ is the dominance coefficient. The fitness of an embryo is multiplicative and is determined by the product of the values of all early-acting loci:

(1 – *d*_e_)*^hme^* (1 – *h*_e_*d*_e_)*^hte^*,

where *hm*_e_ and *ht*_e_ are the numbers of homozygotes and heterozygotes, respectively.

#### Development into seeds

Because of resource limitation, each parent can produce at most *n*_s_ seeds, and the selective abortion of embryos may therefore occur. I assume that embryos with fitness values higher than a threshold of *d*_t_ can potentially develop into seeds. If the number of these embryos is greater than *n*_s_, then *n*_s_ embryos are randomly selected by weighting their fitness values determined by the early-acting loci. Otherwise, all embryos develop into seeds (the number of seeds produced may be smaller than *n*_s_).

#### Late selection

Deleterious effects are assumed to be equal for all mutations occurring at any late-acting locus. The relative fitness values of homozygotes and heterozygotes at a locus are 1 – *d*_l_ (0 ≤ *d*_l_ ≤ 1) and 1 – *h*_l_*d*_l_ (0 ≤ *h*_l_ ≤ 1), where *d*_l_ is the selection coefficient, and *h*_l_ is the dominance coefficient. The fitness of an embryo (seed) during the late selection stage is multiplicative and is determined by the product of the fitness values of all late-acting loci:

(1 – *d*_l_)*^hm^*^l^ (1 – *h*_l_*d*_l_)*^ht^*^l^,

where *hm*_l_ and *ht*_l_ are the numbers of homozygotes and heterozygotes, respectively.

#### Competition for the next generation

Among the seeds produced by the parents in each population, *n* seeds develop into the parents of the next generation. These seeds are randomly selected by weighting their fitness values due to selection caused by late-acting loci.

### Inbreeding depression values

I calculated the inbreeding depression values for each of the final 10 generations of the simulations as follows: I calculated the numbers of seeds produced and the fitness values due to late-acting loci of each seed for each parent, for seeds produced by complete selfing, and for seeds produced by complete outcrossing. Then, I calculated the three means across all parents and obtained three inbreeding depression estimates: 1) one for the number of seeds produced (reflecting selection on early-acting loci), 2) one for embryo survival ability (also reflecting selection on early-acting loci), and 3) one for seed competitive ability (reflecting selection on late-acting loci). These values were averaged over the last 10 generations.

### Simulations carried out

I carried out three kinds of simulations: simulation 1, which includes simulation 1_E_, simulation 1_L_, and simulation 1_EL_; simulation 2, which includes simulation 2_E_, simulation 2_L_, and simulation 2_EL_; and simulation 3, which includes simulation 3_E_, simulation 3_L_, and simulation 3_EL_ (Table 2). Simulation 1 distinguishes the effects caused by the coupling of early- and late-acting mutations from those caused by selective interference. Simulation 2 examines the effects of coupling of early- and late-acting mutations by equalizing the potential total effects of early-acting mutations and those of late-acting mutations among the simulation runs (i.e., the numbers of early-acting loci and late-acting loci are the same among the simulation runs). Simulation 3 examines the effects of the relative numbers of early- and late-acting loci.

**Table 2.** Simulations carried out.

|  | Early-acting loci |  | Late-acting loci |  | Note | Purpose |
| --- | --- | --- | --- | --- | --- | --- |
|  | Number | Occurrence<br>of mutations<br>(indicated by<br>“yes”) | Number | Occurrence of<br>mutations<br>(indicated by<br>“yes”) |  |  |
| Simulation 1 |  |  |  |  | Total number | Comparisons |
| Simulation 1<sub>EL</sub> | 1,000 | Yes | 9,000 | Yes | of loci in | between the |
| Simulation 1<sub>E</sub> | 10,000 | Yes | 0 |  | which | coupling |
| Simulation 1<sub>L</sub> | 0 |  | 10,000 | Yes | mutations may | effects and |
|  |  |  |  |  | occur is the | selective |
|  |  |  |  |  | same. | interference. |
| Simulation 2 |  |  |  |  | Numbers of | Removal of the |
| Simulation 2<sub>EL</sub> | 1,000 | Yes | 9,000 | Yes | early-acting | possible effects |
| Simulation 2<sub>E</sub> | 1,000 | Yes | 9,000 |  | loci and | of having |
| Simulation 2<sub>L</sub> | 1,000 |  | 9,000 | Yes | late-acting | different |
|  |  |  |  |  | loci are the | numbers of |
|  |  |  |  |  | same among | respective loci. |
|  |  |  |  |  | the simulation |  |
|  |  |  |  |  | runs. |  |
| Simulation 3 |  |  |  |  | Numbers of | Examination of |
| Simulation 3<sub>EL</sub> | 9,000 | Yes | 1,000 | Yes | early- and | the effects of |
| Simulation 3<sub>E</sub> | 9,000 | Yes | 1,000 |  | late-acting | the relative |
| Simulation 3<sub>L</sub> | 9,000 |  | 1,000 | Yes | loci are | numbers of |
|  |  |  |  |  | reversed from | early- and |
|  |  |  |  |  | those in | late-acting loci. |
|  |  |  |  |  | simulation 2. |  |

#### Simulation 1: comparisons between the coupling effects and selective interference

For this purpose, comparisons were performed between those simulations in which both coupling effects and selective interference can occur and those in which only selective interference can occur (it is difficult to conduct simulations in which only coupling effects occur). Here, the number of loci in which mutations are allowed to occur and the effects of individual mutations were equalized among the simulations because a higher number of mutations and/or larger effects of individual mutations lead to greater effects of selective interference. Hence, I carried out simulations assuming that only early-acting loci existed (simulation 1_E_), only late-acting loci existed (simulation 1_L_), or both early- and late-acting loci existed (simulation 1_EL_). I made the following assumptions for these simulations:

1. In all of these simulations, the total number of loci is 10,000; the numbers of early- and late-acting loci are, respectively, 10,000 and 0 in simulation 1_E_, 0 and 10,000 in simulation 1_L_, and 1,000 and 9,000 in simulation 1_EL_.
2. The selection coefficient is the same for all mutations (*d*_e_ = *d*_l_ = 0.05, 0.2, or 0.5).
3. The dominance coefficient is also the same for all mutations (*h*_e_ = *h*_l_ = 0.02).
4. There is no threshold for the development into seeds (i.e., *d*_t_ = 0), and hence, the form of selection is the same for embryo competition, where early-acting genes express, and for seed competition for the next generation, where late-acting genes express.
5. The strength of selection is the same for embryo competition and seed competition; the numbers of embryos and seeds produced by a parent, *n*_o_ and *n*_p_, are 36 and 6, respectively. Thus, 1/6 of embryos or seeds are selected in both the embryo and seed competitions.

The potential effects of mutations are the same in all of these simulations, and selective interference can take place, but the coupling of early- and late-acting mutations does not occur in simulations 1_E_ and 1_L_.

Embryos developing into seeds are randomly selected in simulation 1_L_, and seeds developing into the next generation are randomly selected in simulation 1_E_. In all of simulation 1, *n*_link_ = 5 and *n*_locus_ = 2,000, and in simulation 1_EL_, each chromosome contains 200 early-acting loci and 1,800 late-acting loci. The other parameter values used were *s* = 0.3–0.8, *m* = 0.00005, *r* = 1, and *n* = 300. The population size remained constant.

#### Simulation 2: comparisons when the numbers of early-acting loci and late-acting loci are the same among the simulation runs

In these simulations, the numbers of early-acting loci and late-acting loci are the same among the simulation runs, in contrast to simulation 1. This equalization removes the possible effects of having different numbers of respective loci on the maintenance of mutations. For example, the greater number of early-acting loci (or late-acting ones) will result in the higher degree of inbreeding depression caused by early-acting loci (or late-acting loci). I hence assumed that, in total, 1,000 early- and 9,000 late-acting loci exist in simulation 2. Then, I carried out simulations in which mutations were allowed only in early-acting loci (simulation 2_E_), only in late-acting loci (simulation 2_L_), or in both the early- and late-acting loci (simulation 2_EL_). The potential effects of mutations caused by early-acting loci are the same between simulations 2_E_ and 2_EL_, and those caused by late-acting loci are also the same between simulations 2_L_ and 2_EL_, where the coupling of early- and late-acting mutations occurs only in simulation 2_EL_.

For the main runs of simulation 2, I assumed *n*_link_ = 5 and *n*_locus_ = 2,000, and for each haplotype genome, one chromosome contains 1,000 early- and 1,000 late-acting loci, and the other chromosomes contain only 2,000 late-acting loci. Other assumptions were also made for the other runs (see below). For the other parameters, I used the following values: *s* = 0.3–0.8; *m* = 0.00005; *d*_e_ = 1; *d*_l_ = 0.05, 0.2, or 0.5; *h*_e_ = *h*_l_ = 0.02; *d*_t_ = 0.2; *r* = 1; *n* = 300; *n*_s_ = 10; and *n*_o_ = *n*_p_ = 20. The population size remained constant.

#### Effects of linkage

In addition, to examine the effects of linkage on the results, I carried out the following two kind of runs for simulation 2: 1) *n*_link_ = 5 and *n*_locus_ = 2,000; for each haplotype genome, each chromosome contains 200 early- and 1,800 late-acting loci (1,000 early- and 9,000 late-acting loci in total, as in the main runs of simulation 2) and 2) all loci are unlinked, all loci segregate independently, and there are 1,000 early- and 9,000 late-acting loci. Simulation 2 was used because if the numbers of early-acting loci and late-acting loci differ among the simulation runs, factors other than the linkage between early- and late-acting loci might affect the results, in which case the effects of the linkage might not be detected. The other parameter values used were the same as those in the main runs of simulation 2, except that *r* was not assumed for the runs without the linkage of chromosomes.

#### Effects of the dominance coefficient

Using simulation 2, I also examined the effects of the dominance coefficient in the late-acting loci, *h*_l_. The mean dominance coefficient is 0.1 for *Caenorhabditis elegans* (PETERS *et al*. 2003), 0.17 for *Drosophila melanogaster* (FRY AND NUZHDIN 2003), and 0.197 for nonlethal mutations in yeasts (SZAFRANIEC *et al*. 2003), with large variances. Hence, I assumed *h*_l_ = 0.02, 0.1, 0.2, and 0.3 for the late-acting loci. More deleterious mutations are more likely to be recessive than less deleterious mutations (HUBER *et al*. 2018), and early-acting mutations are highly deleterious and nearly recessive (see Introduction of PORCHER AND LANDE 2005b). Hence, I assumed *h*_e_ = 0.02 for the early-acting loci for all simulations. Simulation 2 was used because if the numbers of early-acting loci and late-acting loci differ among the simulation runs, factors other than the dominance coefficient might affect the results. The other parameter values used were the same as those in the main runs of simulation 2.

#### Simulation 3: comparisons when the number of early-acting loci is greater than that of late-acting loci

I also examine the effects of the relative numbers of early- and late-acting loci by exchanging the numbers of respective loci used in simulation 2. In this simulation, *n*_link_ = 5 and *n*_locus_ = 2,000, as in simulation 2, but for each haplotype genome, each chromosome contained 1,800 early-acting loci and 200 late-acting loci. Hence, the effects of selective interference could be strong in the early-acting loci in this simulation, whereas those effects could be strong in the late-acting loci in simulation 2. The other parameter values used were the same as those in the main runs of simulation 2.

## Results

Figure 1 shows examples of the dynamics of the deleterious, recessive mutations that occur in the early- and late-acting loci (simulation 2; 1,000 and 9,000 early- and late-acting loci, respectively). The simulations start with no mutations at either the early- or late-acting loci. If mutations are allowed to occur in both sets of loci, then their numbers increase over the generations in both loci (solid lines). However, if mutations are allowed to occur in only the early- or late-acting loci (1,000 and 9,000 locus mutations, respectively), then they remain rare in both sets of loci (dotted lines).

**Figure 1.**
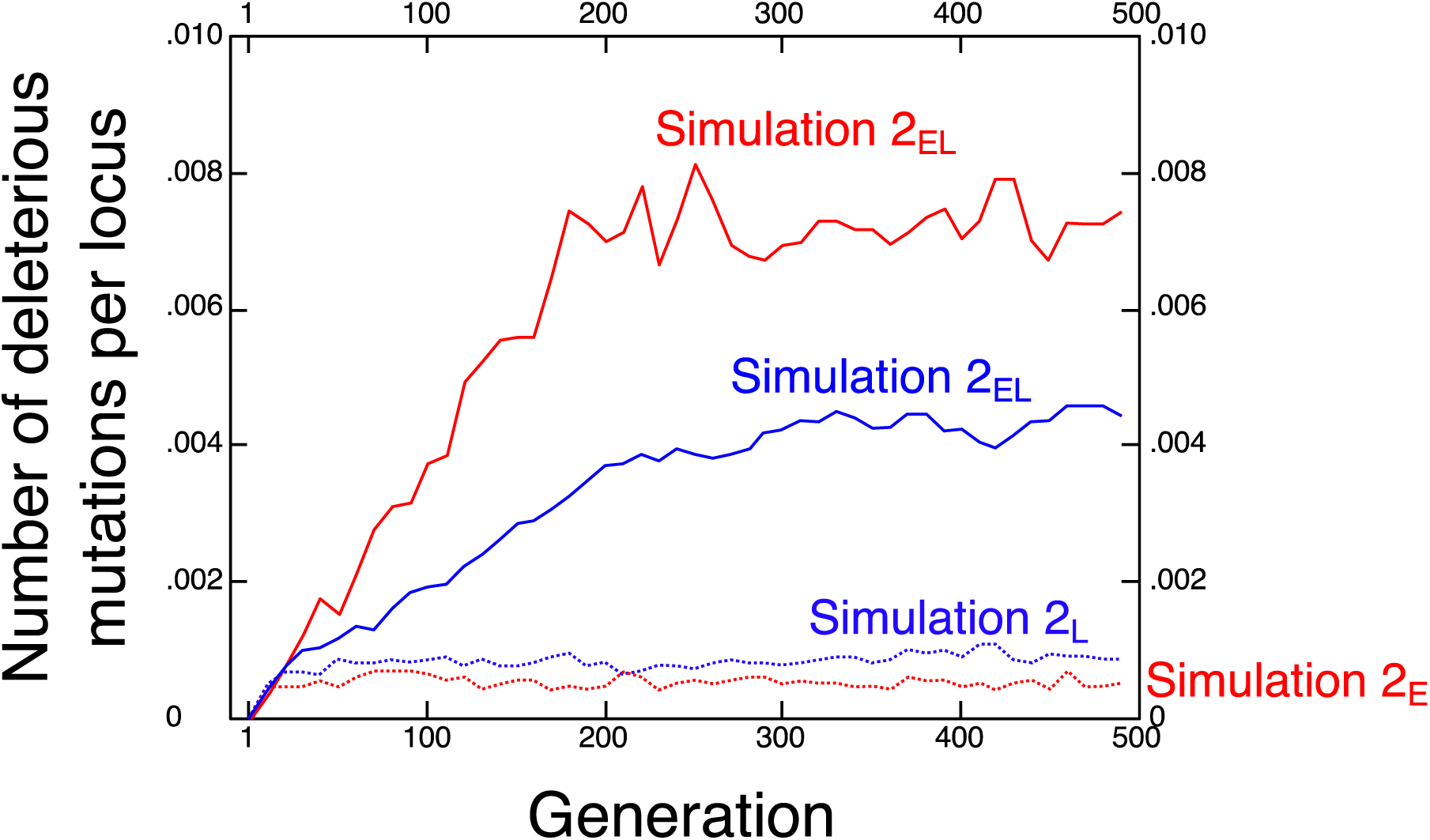
Examples of the dynamics of deleterious, recessive mutations occurring in early- and late-acting loci (simulation 2). Red lines; early-acting mutations, blue lines; late-acting mutations, solid lines; mutations occur in both sets of loci (simulation 2_EL_), dotted lines; mutations occur in the loci of the corresponding color but not in the other set of loci (simulations 2_E_ and 2_L_).

### Simulation 1

Under most parameter values, the numbers of early- and late-acting mutations (left and right panels, respectively, in Fig. 2) that are maintained are greater if both early- and late-acting loci exist (red circles; simulation 1_EL_) than if only early-acting loci (gray circles; simulation 1_E_) or only late-acting loci exist (black circles; simulation 1_L_). Thus, the presence of both sets of loci promotes the maintenance of mutations under the same effects of selective interference. Here, the differences are smaller for late-acting mutations. This trend may be because the numbers of late-acting loci are similar between simulations 1_EL_ and 1_L_ (9,000 and 10,000, respectively) and, hence, the effects of selective interference should be similar for late-acting loci (also see the explanation for the maintenance of mutations in simulations 2 and 3). On the other hand, if *s* = 0.7 - 0.8 and *d*_e_ = *d*_l_ = 0.2 or = 0.5, the numbers of early-acting mutations that are maintained are lower if both sets of loci exist than if only early-acting loci exist. Additionally, if *s* is low or high, the numbers of late-acting mutations that are maintained are similar between the simulations when both sets of loci exist and when only late-acting loci exist.

**Figure 2.**
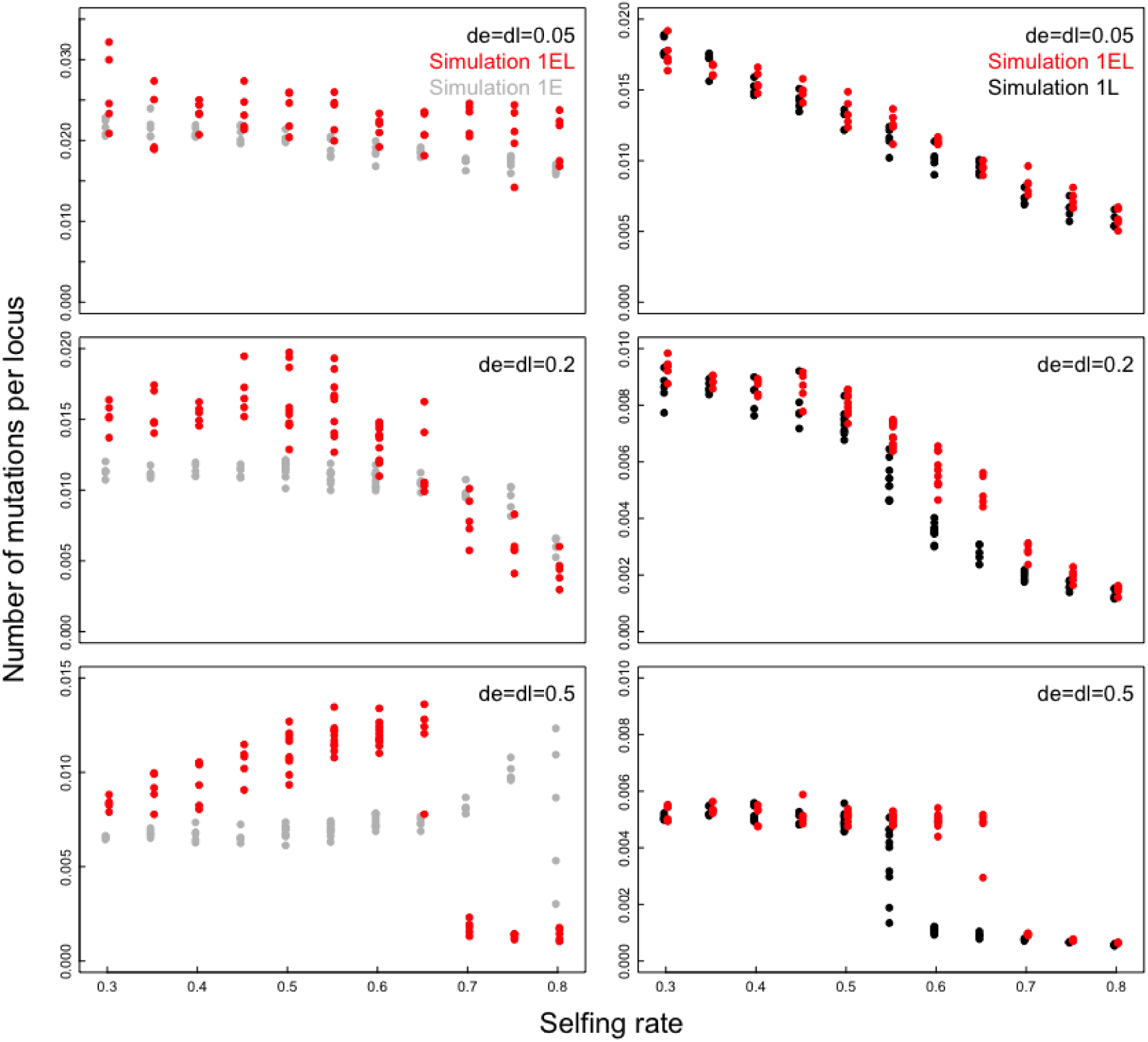
Comparisons of the numbers of mutations maintained between the coupling effects and selective interference (simulation 1). The dependences of the number of early-acting mutations (left panels) and late-acting mutations (right panels) per locus after 500 generations on the selfing rate, *s*, with the selection coefficients against an individual early- or late-acting mutation, *d*_e_ or *d*_l_, shown. The total number of loci in which mutations are allowed to occur is 10,000 (*n*_link_ = 5 and *n*_locus_ = 2,000) in all simulations. Red circles; simulation 1_EL_, in which the numbers of early-acting and late-acting loci are 1,000 and 9,000, respectively (with each chromosome containing 200 early-acting loci and 1,800 late-acting loci), and mutations occur in both the early- and late-acting loci. Gray circles; simulation 1_E_, in which the numbers of early-acting and late-acting loci are 10,000 and 0, respectively, and mutations occur only in the early-acting loci. Black circles; simulation 1_L_, in which the numbers of early-acting and late-acting loci are 0 and 10,000, respectively, and mutations occur only in the late-acting loci. The simulation results of 5 or 10 runs are shown for each combination of parameter values.

### Simulations 2 and 3

#### Maintenance of mutations

The numbers of early-acting mutations (left panels in Fig. 3) that are maintained are greater if mutations occur in both the early- and late-acting loci (red circles; simulation 2_EL_). Early-acting mutations are not maintained only if *s* is very high. In contrast, if mutations do not occur in the late-acting loci, then the numbers of early-acting mutations that are maintained are very low, even if mutations occur in the early-acting loci (gray circles in Fig. 3; simulation 2_E_).

**Figure 3.**
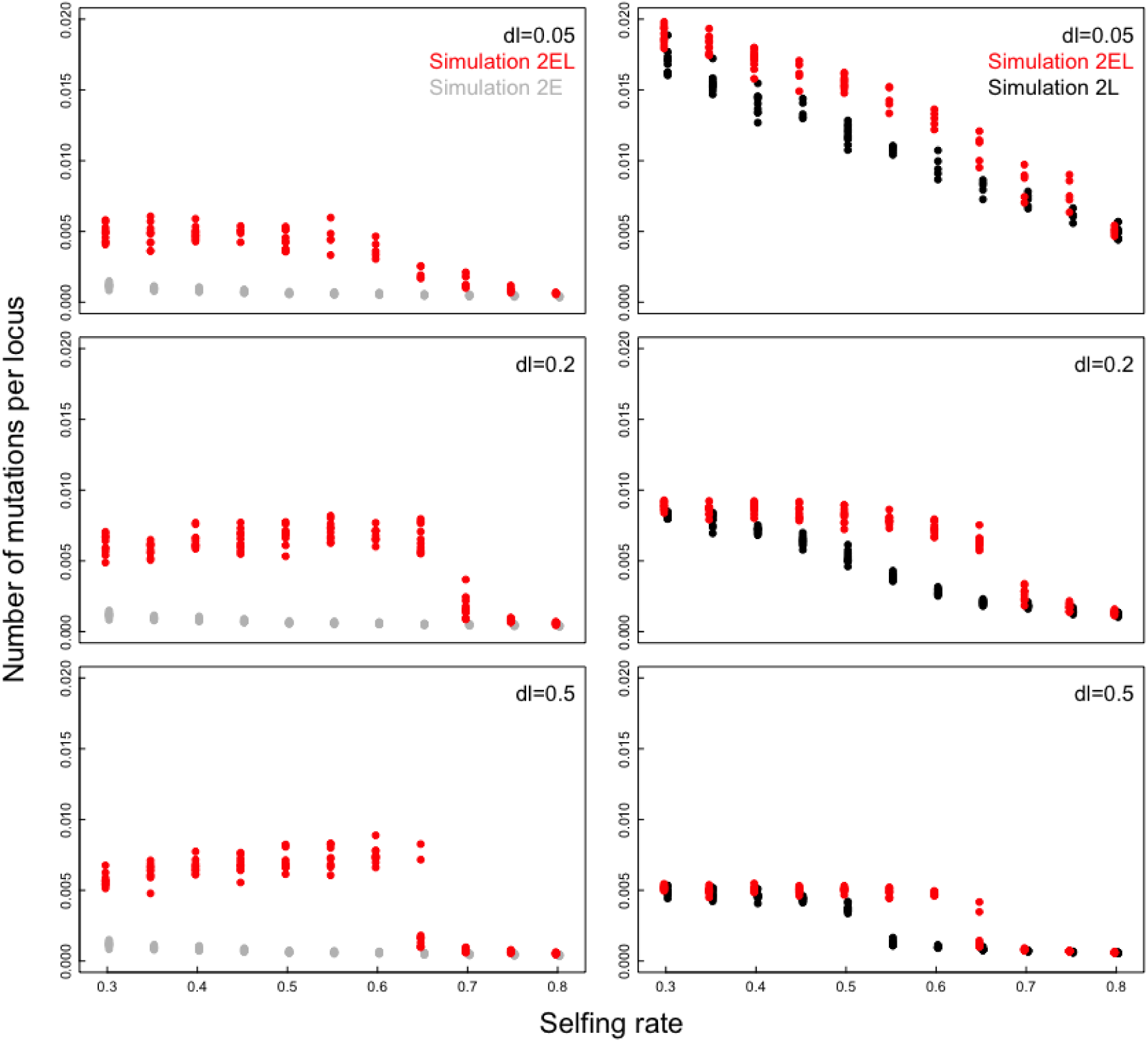
Comparisons of the numbers of mutations maintained among the simulation runs having the same number of early-acting loci and the same number of late-acting loci (simulation 2). The dependences of the number of mutations per locus after 500 generations on the selfing rate, *s*, with the selection coefficient against an individual late-acting mutation, *d*_l_, shown. In all panels, the numbers of early-acting and late-acting loci are 1,000 and 9,000, respectively, and the early-acting mutations are lethal (*d*_e_ = 1) if they occur. Red circles; simulation 2_EL_, in which mutations occur in both the early- and late-acting loci. Gray circles; simulation 2_E_, in which mutations occur in the early-acting loci but not in the late-acting loci. Black circles; simulation 2_L_, in which mutations occur in the late-acting loci but not in the early-acting loci. The simulation results of 5 or 10 runs are shown for each combination of parameter values.

The number of late-acting mutations (right panels in Fig. 3) is higher when mutations occur in both the early- and late-acting loci (red circles; simulation 2_EL_), particularly if *s* is moderate. Late-acting mutations are rare only if *s* is very high. However, even if mutations do not occur in the early-acting loci, the numbers of late-acting mutations that are maintained tend to be high (black circles; simulation 2_L_). This trend may be due to selective interference, which promotes their maintenance under the assumption that the number of late-acting loci (9,000) is greater than that of early-acting loci (1,000). In fact, if the numbers of early-and late-acting loci are 9,000 and 1,000, respectively, then the opposite results are observed (Fig. S1; simulation 3). The numbers of late-acting mutations that are maintained are very small if the mutations do not occur in the early-acting loci, whereas the numbers of early-acting mutations that are maintained are high even if mutations do not occur in the late-acting loci.

The mechanism modeled here increases the frequency of late-acting mutations that are present as heterozygotes; more late-acting mutations are present as heterozygotes if mutations occur in both the early- and late-acting loci than if they occur only in the late-acting loci (Fig. S2). This case obtains particularly if *s* is not low and *d*_l_ is low.

#### Inbreeding depression

The inbreeding depression values related to the number of seeds produced (left panels in Fig. 4) and embryo survival ability (Fig. S3) were also greater if mutations occurred in both the early- and late-acting loci, except for very high values of *s*. The inbreeding depression values related to seed competitive ability (right panels in Fig. 4) also show similar patterns to the number of mutations maintained (Fig. 3).

**Figure 4.**
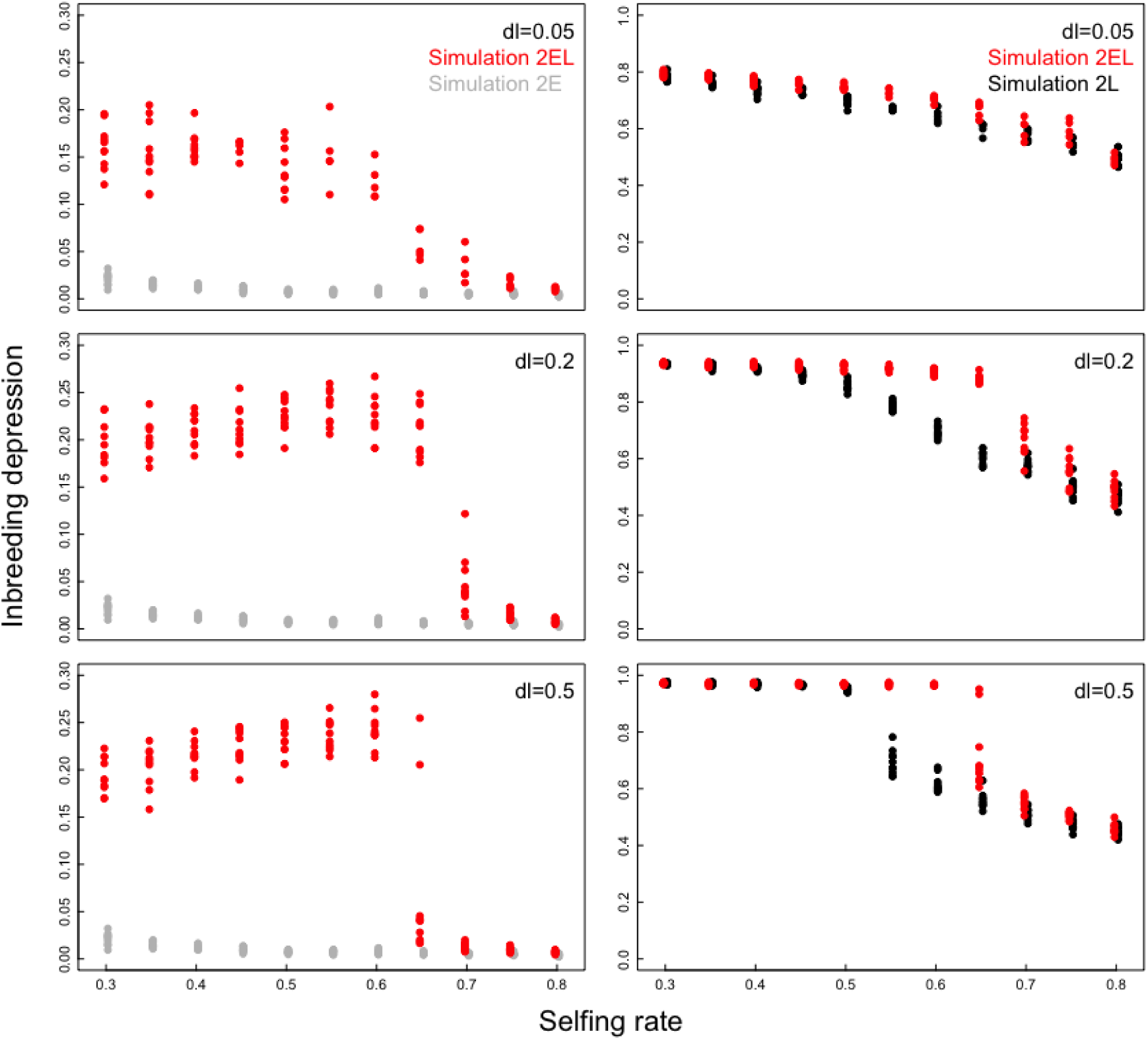
Comparisons of the degree of inbreeding depression among the simulation runs having the same number of early-acting loci and the same number of late-acting loci (simulation 2). The dependences of the effects of inbreeding depression on seed production (left panels) or seed competitive ability (right panels) after 500 generations on the selfing rate, *s*, with the selection coefficient against an individual late-acting mutation, *d*_l_, shown. In all panels, the numbers of early-acting and late-acting loci are 1,000 and 9,000, respectively, and the early-acting mutations are lethal (*d*_e_ = 1) if they occur. Red circles; simulation 2_EL_, in which mutations occur in both the early- and late-acting loci. Gray circles; simulation 2_E_, in which mutations occur in the early-acting loci but not in the late-acting loci. Black circles; simulation 2_L_, in which mutations occur in the late-acting loci but not in the early-acting loci. The simulation results of 10 runs are shown for each combination of parameter values.

#### Effects of the overproduction of ovules

The extent to which ovules are overproduced affects the preservation of deleterious mutations and thus the genetic load maintained in the population (Fig. 5). In this figure, the number of ovules produced by a plant, *n*_o_, is equal to 20 in all panels; therefore, a greater value of *n*_s_ indicates a lower degree of ovule overproduction. If the selfing rate is high (*s* = 0.6, 0.65, and 0.7 in Fig. 5), a small or moderate degree of overproduction is sufficient for maintenance, and if the selfing rate is not very high (*s* = 0.5 in Fig. 7), deleterious mutations are maintained without overproduction if both early- and late-acting mutations occur (red circles; simulation 2_EL_). However, if mutations do not occur either in the late- or early-acting loci (gray or black circles; simulation 2_E_ or 2_L_), the number of mutations maintained is low and is almost independent of the maximum number of seeds produced.

**Figure 5.**
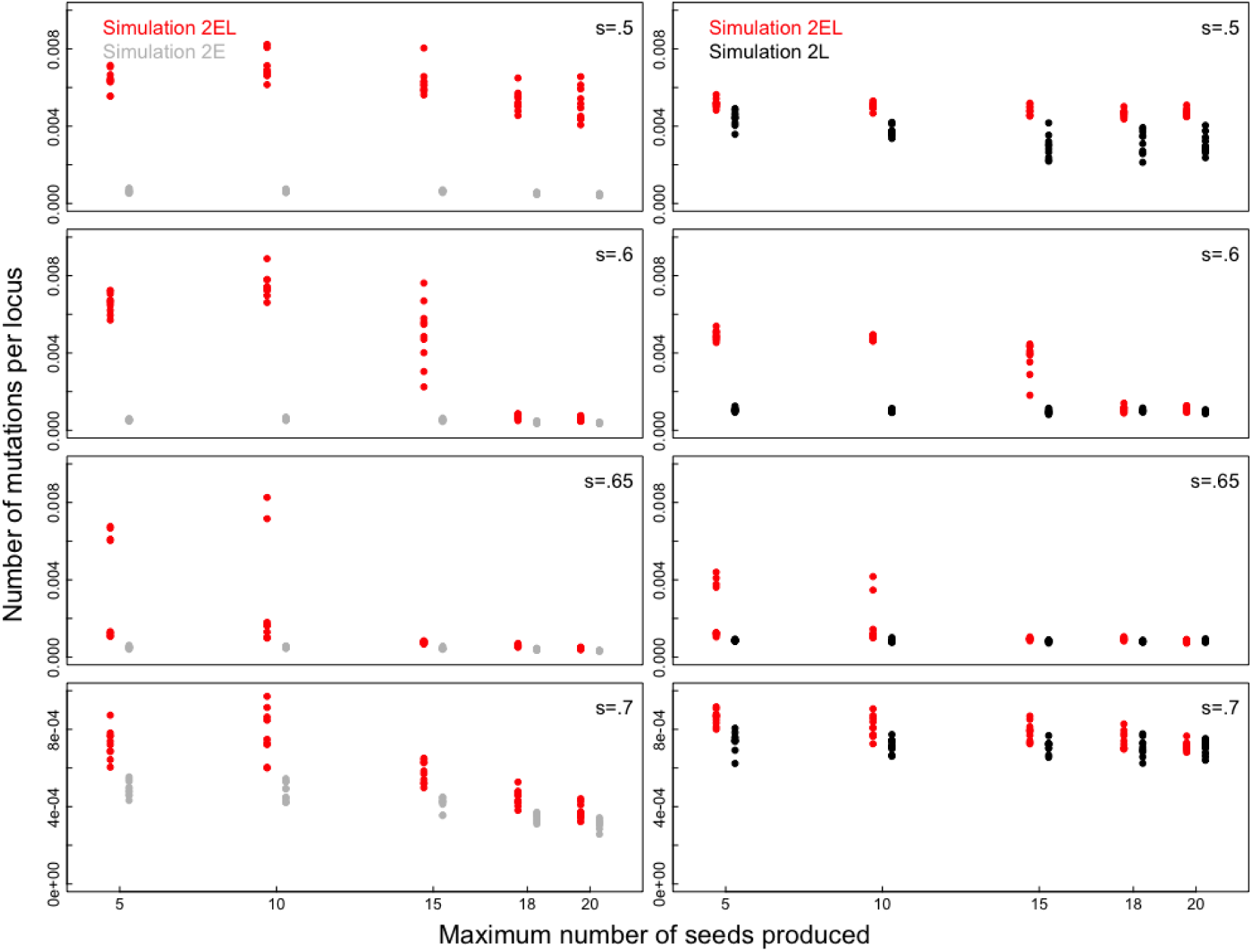
The effects of the degree of overproduction of ovules (simulation 2). The dependences of the number of early-acting mutations (left panels) and late-acting mutations (right panels) per locus after 500 generations on the selfing rate, *s*, with the maximum number of seeds produced by a plant, *n*_s_, shown. The number of ovules produced by a plant, *n*_o_, is equal to 20 in all panels; therefore, a greater value of *n*_s_ indicates a lower degree of ovule overproduction. The selection coefficients against an individual early-acting mutation and an individual late-acting mutation are *d*_e_ = 1 and *d*_l_ = 0.5, respectively, in all panels. Red circles; simulation 2_EL_, in which mutations occur in both the early- and late-acting loci. Gray circles; simulation 2_E_, in which mutations occur in the early-acting loci but not in the late-acting loci. Black circles; simulation 2_L_, in which mutations occur in the late-acting loci but not in the early-acting loci. The simulation results of 10 runs are shown for each combination of parameter values.

#### Effects of linkage

Linkage does not qualitatively affect the results. The results are very similar to those presented in Figs. 3 and 4, even if each chromosome contains 200 early-acting loci and 1,800 late-acting loci (*n*_link_ = 5 and *n*_locus_ = 2000; Fig. S4) or all loci are unlinked and segregate independently (Fig. S5). However, the degree of inbreeding depression in seed production becomes high (relative to that presented in Fig. 4) when both early- and late-acting mutations occur.

#### Effects of the dominance coefficient

Irrespective of the value of the dominance coefficient in the late-acting loci, *h*_l_, the numbers of early-acting mutations that are maintained (Fig. S6) and the inbreeding depression values (Figs. S7 and S8) are higher if mutations occur in both the early- and late-acting loci than if mutations do not occur in the late-acting loci, although the differences between these two conditions become small with increases in *h*_l_. On the other hand, the numbers of late-acting mutations that are maintained (Fig. S9) and the inbreeding depression values (Figs. S10) tend to be slightly higher if mutations occur in both the early- and late-acting loci than if mutations do not occur in the early-acting loci, but the differences are very small. This situation may also occur because selective interference promotes their maintenance under the assumption that the number of late-acting loci (9,000) is great.

## Discussion

### Coupling effects of early- and late-acting mutations

Early- and late-acting mutations are maintained if mutations occur in both sets of loci but are not maintained if they occur only in either the early- or late-acting loci (Figs. 1-4). Thus, for maintenance of mutations to occur, mutations must take place in both loci, which implies that the occurrence of early-acting mutations promotes the spread of late-acting mutations and vice versa.

These results can be understood intuitively. Let *E* and *L* be early-acting and late-acting deleterious alleles, respectively, and *e* and *l* be early- and late-acting wild-type alleles, respectively. Consider embryo production by *EeLl* parents, and assume that an *EeLl* parent happened to produce *EELL* embryos at a higher frequency and *EELl* and *EeLL* embryos at low frequencies. For example,

parent A; *EELL*, *EeLl*, *Eell*, *eeLl*, *eell*

parent B; *EELl*, *EeLL*, *Eell*, *eeLl*, *eell*

The embryos of parent A are more successful than those of parent B because only *EELL* embryos are unable to survive to the next generation in parent A, whereas both *EELl* and *EeLL* embryos are unable to survive in parent B. In addition, because the frequencies of *E* and *L* alleles in the other embryos are higher in parent A than in parent B, more *E* and *L* alleles are passed to the next generation by parent A. Thus, for given frequencies of *EE* and *LL* in the embryos produced by a parent, more *E* and *L* alleles are passed to the next generation if *EELL* embryos happened to be produced more frequently, such that *EELl* and *EeLL* embryos are produced at lower frequencies. On the other hand, if embryos are overproduced, parent A is not disadvantageous compared to the parent with genotypes such as *eell* because a similar number of seeds develop in both parents. Thus, in comparison to other parents, parent A produces successful seeds bearing heterozygous early- and late-acting mutations at higher frequencies. Mutations spread through these heterozygotes.

### Differences from selective interference

The coupling effect of early- and late-acting mutations differs from selective interference and, compared to selective interference, more strongly promotes the maintenance of mutations (Fig. 2). In these simulations, the total number of loci in which mutations are allowed to occur and the form and strength of selection are the same. However, the numbers of mutations that are maintained are much greater when coupling occurs than when selective interference alone occurs for most of the examined parameters. Embryo competition, where early-acting genes express, is a form of within-plant competition, whereas seed competition, where late-acting genes express, is a form of among-plant competition. Thus, the effects of selective interference can be expected to differ between embryo competition and seed competition. Accordingly, selective interference should lead to the maintenance of the greatest number of mutations when only early- or late-acting genes exist. However, the number of mutations maintained is greatest when both genes exist, indicating that the presence of both mutations is important.

The difference in the number of mutations maintained between the coupling effect and selective interference may be due to differences between these phenomena in the mechanisms by which mutations are maintained. In selective interference, mutations spread through outcrossed embryos, and hence, strong selection against selfed embryos is necessary; a lower survival rate of selfed embryos leads to a greater number of maintained mutations. On the other hand, in the coupling effect, as explained above, mutations spread through selfed embryos as well as through outcrossed ones through the coupling of early- and late-acting mutations. Because the contribution of outcrossed embryos should be the same, the coupling effect promotes the maintenance of mutations more strongly than does selective interference.

The exception is the maintenance of early-acting loci when selfing rates and selection coefficients are very high; more early-acting mutations are maintained when there are 10,000 early-acting loci (Fig. 2). In this case, very strong inbreeding depression occurs, and almost all selfed zygotes die, resulting in very strong selective interference. However, it seems unlikely that mutations are maintained by selective interference when selfing rates and selection coefficients are very high. In the simulations, 10,000 early-acting loci exist. (This parameter value was used for comparisons between the coupling effect and selective interference rather than to derive realistic predictions.) Given that the total numbers of loci are 25,498 in *Arabidopsis thaliana* (KAUL *et al*. 2000) and approximately 35,000 in tomato (SATO *et al*. 2012), it is doubtful that 10,000 early-acting loci actually exist. Thus, very strong mutation effects caused by a very large number of early-acting loci and very deleterious mutations might have promoted the strong purging of selfed embryos to an unrealistic extent.

### Effects of the overproduction of ovules

An increase in the overproduction of ovules results in a higher number of deleterious mutations that are maintained (Fig. 5). Porcher and Lande (2005b) similarly showed that a higher degree of overproduction leads to the maintenance of a greater number of mutations. Several adaptive reasons may explain the overproduction of ovules (summarized by PORCHER AND LANDE 2005b) [e.g., enhanced male reproductive success via increased pollen production and export generated from excess flowers (SUTHERLAND 1987; BURD AND CALLAHAN 2000) and enhanced female reproductive success via pollinator attraction (BURD 1998)]. In any case, the present model is applicable to most plant species because the selective abortion of embryos may occur, as not all embryos develop into seeds. Hence, given that overproduction of ovules is very common (LEE 1988), the present model may apply to most plant species.

However, the overproduction of ovules alone cannot lead to the maintenance of deleterious mutations; the coupling of early- and late-acting mutations is necessary. This is because if mutations do not occur either in the late- or early-acting loci, the number of mutations maintained is low and is almost independent of the degree of overproduction (gray and black circles in Fig. 5; simulations 2_E_ and 2_L_). In contrast to the model of Porcher and Lande (2005b), in which the same mutations become both early acting and late acting, the present model distinguishes early- and late-acting loci, and the effects of early- and late-acting mutations appear during their respective stages. Then, if mutations do not occur in the early-acting loci, embryos developing into seeds are randomly selected, and the overproduction of ovules does not affect the relative frequencies of homozygous and heterozygous late-acting mutations and hence does not affect the maintenance of those mutations. Furthermore, if mutations do not occur in the late-acting loci, the function of overproduction—embryos that will not become vigorous seeds are removed—does not exist. A high degree of overproduction rather enhances the removal of early-acting mutations because of the severe competition among embryos and hence does not contribute to the maintenance of early-acting mutations.

### Effects of linkage

Linkage is not an important factor in maintenance of mutations, as evidenced by the similar results among the simulations in which linkage exists (Figs. 3 and 4), all loci are unlinked, or all loci segregate independently (Fig. S5). Thus, the present results were not caused by linkage disequilibrium.

### Effects of the dominance coefficient

The coupling effect occurs even if the dominance coefficient in the late-acting loci, *h*_l_, is higher than 0.02, although the effect diminishes with increasing dominance coefficient (Figs. S6-S10). In organisms, mean values of the dominance coefficient range between approximately 0.1 and 0.2, but the values are highly variable among mutations (FRY AND NUZHDIN 2003; PETERS *et al*. 2003; SZAFRANIEC *et al*. 2003; HUBER *et al*. 2018). Hence, many mutations with dominance coefficients much smaller than those means are included, and such mutations should be more effectively maintained by the coupling effect than mutations with high dominance coefficients. Thus, mutations with small dominance coefficients may contribute more strongly than those with large dominance coefficients to the high inbreeding depression in selfing populations.

### Implications for studies of selfing and inbreeding depression

The present results can aid in the understanding of empirical studies of selfing and inbreeding depression. Here, I discuss several topics.

The mechanism modeled here increases the frequency of late-acting mutations present as heterozygotes (Fig. S2). If late-acting mutations are present as heterozygotes, seeds are more vigorous than they are if the same number of mutations are maintained at higher homozygote frequencies. Hence, the risk of extinction of populations caused by the accumulation of deleterious mutations may be lower if mutations are present as heterozygotes in the seeds. Thus, the probability that populations with high selfing rates can persist is higher under the present mechanism than under selective interference alone.

The present study has implications for the study of reproductive allocation in flowers, i.e., the allocation to pollinator attraction and male and female organs, as it shows that the overproduction of ovules affects the extent of inbreeding depression and, hence, the vigor of selfed seeds. Thus, ovule production and outcrossing strategy may be related to each other in a previously unconsidered way. For example, a change in the number of ovules produced results in a change in the vigor of selfed seeds, leading to changes in the evolutionarily stable selfing rate and resource allocation to enhance outcrossing. The adaptive significance of ovule overproduction, such as through bet-hedging and selective abortion (KOZLOWSKI AND STEARNS 1989), may also change under the present model. Thus, the present study may contribute to future studies on these and other topics.

## Acknowledgements

This study was supported in part by a grant-in-aid from the Japanese Ministry of Education, Culture, Sports, Science and Technology.

## Supplemental Material

**Figure S1.**
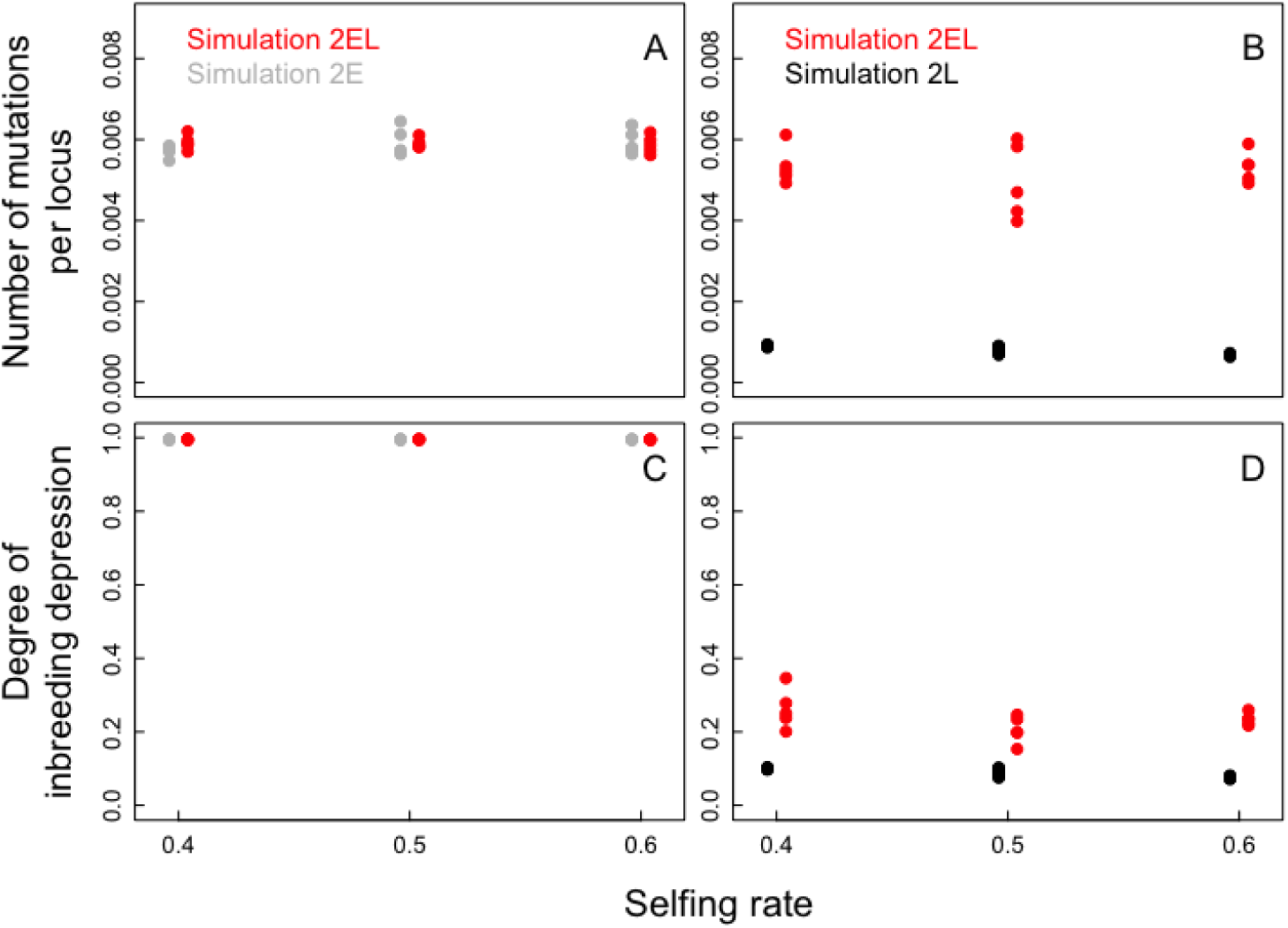
The effects of the relative numbers of early-acting and late-acting loci (simulation 3): comparison with the results in which each chromosome contains 1,800 early-acting loci and 200 late-acting loci (*n*_link_ = 5 and *n*_locus_ = 2000). The dependences of the numbers of early- (A) and late-acting (B) mutations per locus after 500 generations and the degree of inbreeding depression related to the number of seeds produced (C) and the seed competitive ability (D) on the selfing rate, *s*, are shown. In all panels*, d*_e_ = 1 and *d*_l_ = 0.5 if early- and late-acting mutations occur, respectively. Red circles; simulation 2_EL_, in which mutations occur in both the early- and late-acting loci. Gray circles; simulation 2_E_, in which mutations occur in the early-acting loci but not in the late-acting loci. Black circles; simulation 2_L_, in which mutations occur in the late-acting loci but not in the early-acting loci. The simulation results of 5 runs are shown for each combination of parameter values.

**Figure S2.**
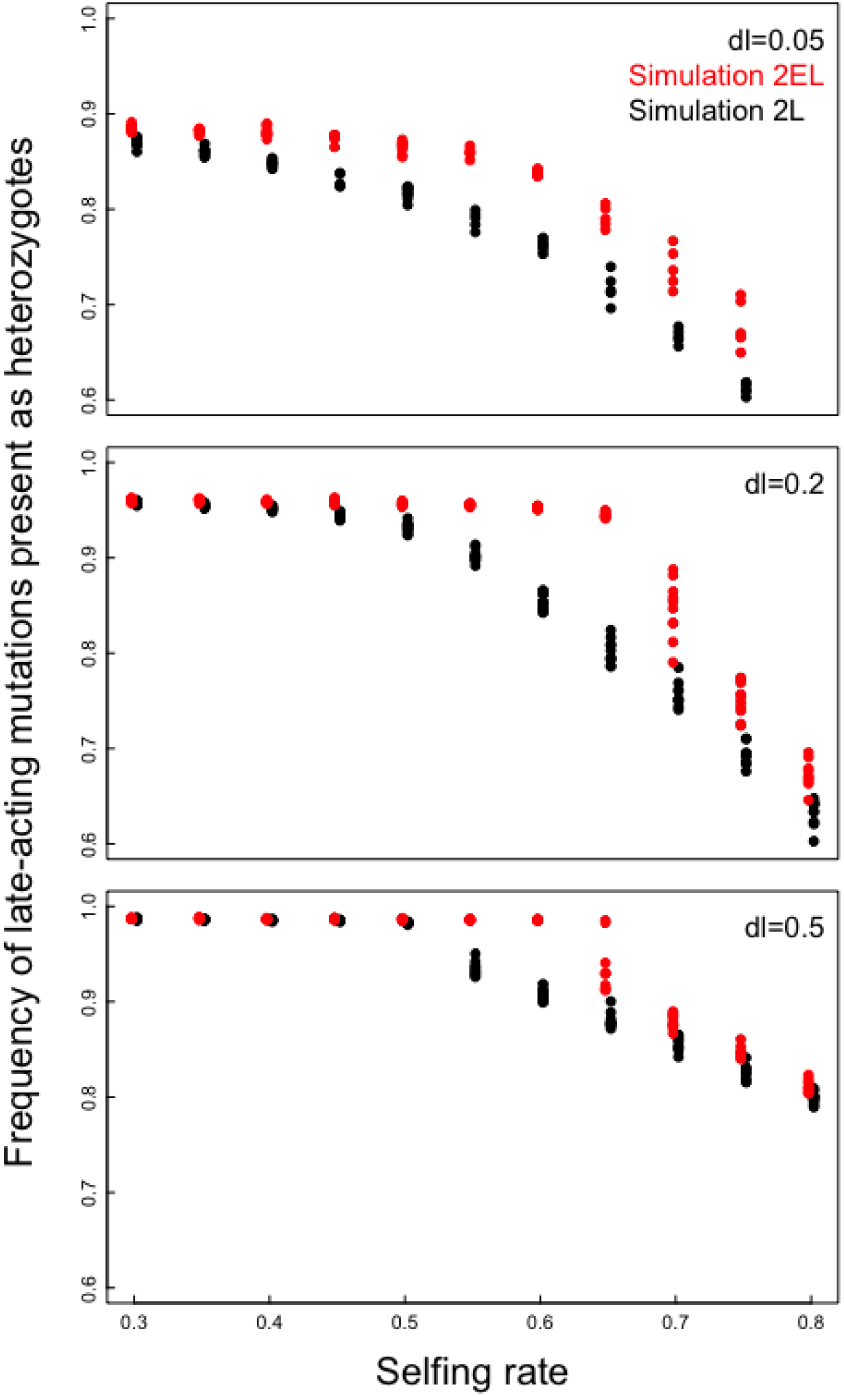
The dependences of the relative frequencies of late-acting mutations present as heterozygotes in the seeds produced after 500 generations on the selfing rate, *s*, and the selection coefficient against an individual late-acting mutation, *d*_l_ (simulation 2). In all panels, the early-acting mutations are lethal (*d*_e_ = 1) if they occur. Red circles; simulation 2_EL_, in which mutations occur in both the early- and late-acting loci. Black circles; simulation 2_L_, in which mutations occur in the late-acting loci but not in the early-acting loci. The simulation results of 10 runs are shown for each combination of parameter values.

**Figure S3.**
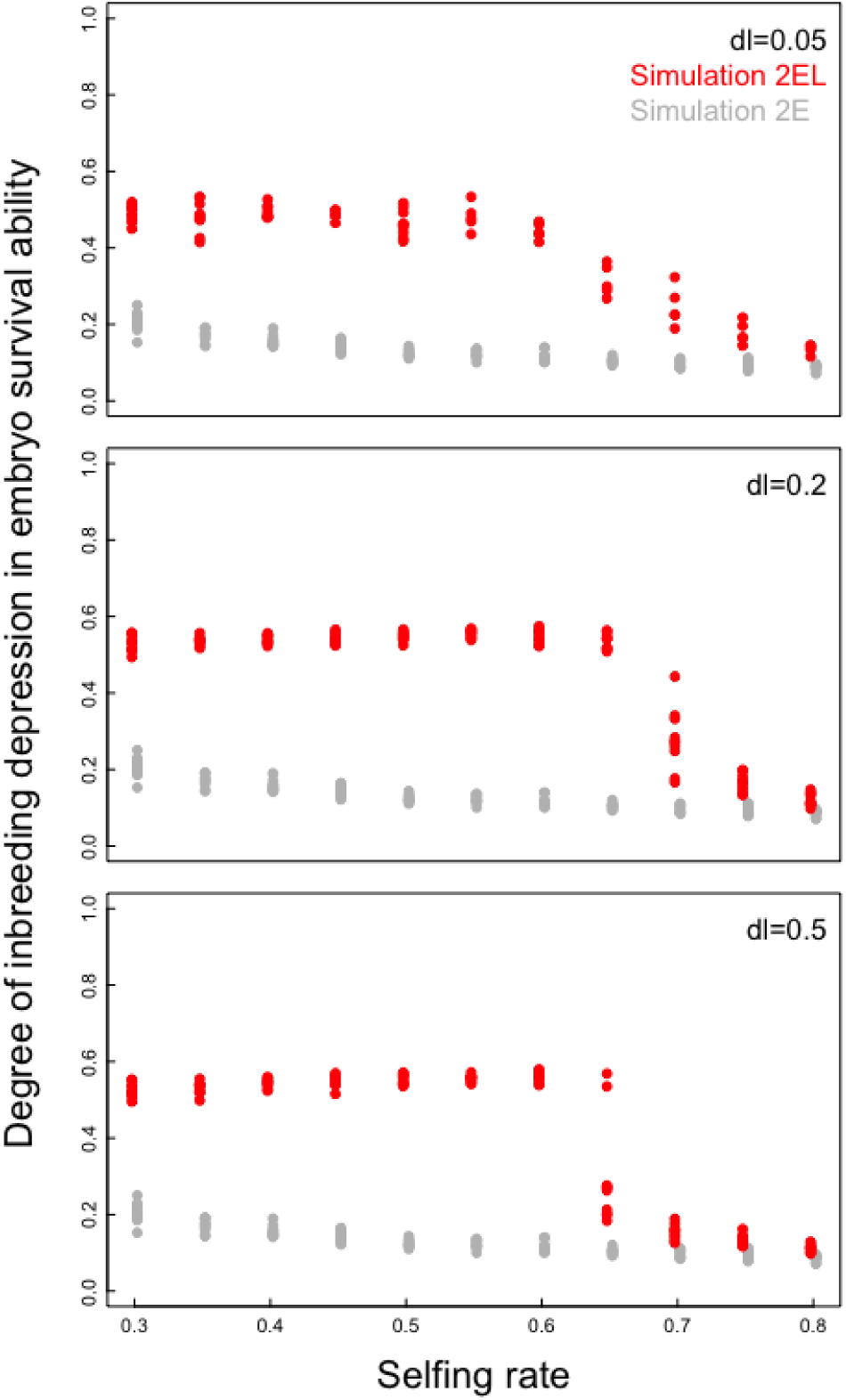
Comparisons of the degree of inbreeding depression among the simulation runs having the same number of early-acting loci and the same number of late-acting loci (simulation 2). The dependences of the effects of inbreeding depression on the embryo survival ability after 500 generations on the selfing rate, *s*, with the selection coefficient against an individual late-acting mutation, *d*_l_, shown. The early-acting mutations are lethal (*d*_e_ = 1) in all panels. Red circles; simulation 2_EL_, in which mutations occur in both the early- and late-acting loci. Gray circles; simulation 2_E_, in which mutations occur in the early-acting loci but not in the late-acting loci. The simulation results of 10 runs are shown for each combination of parameter values.

**Figure S4.**
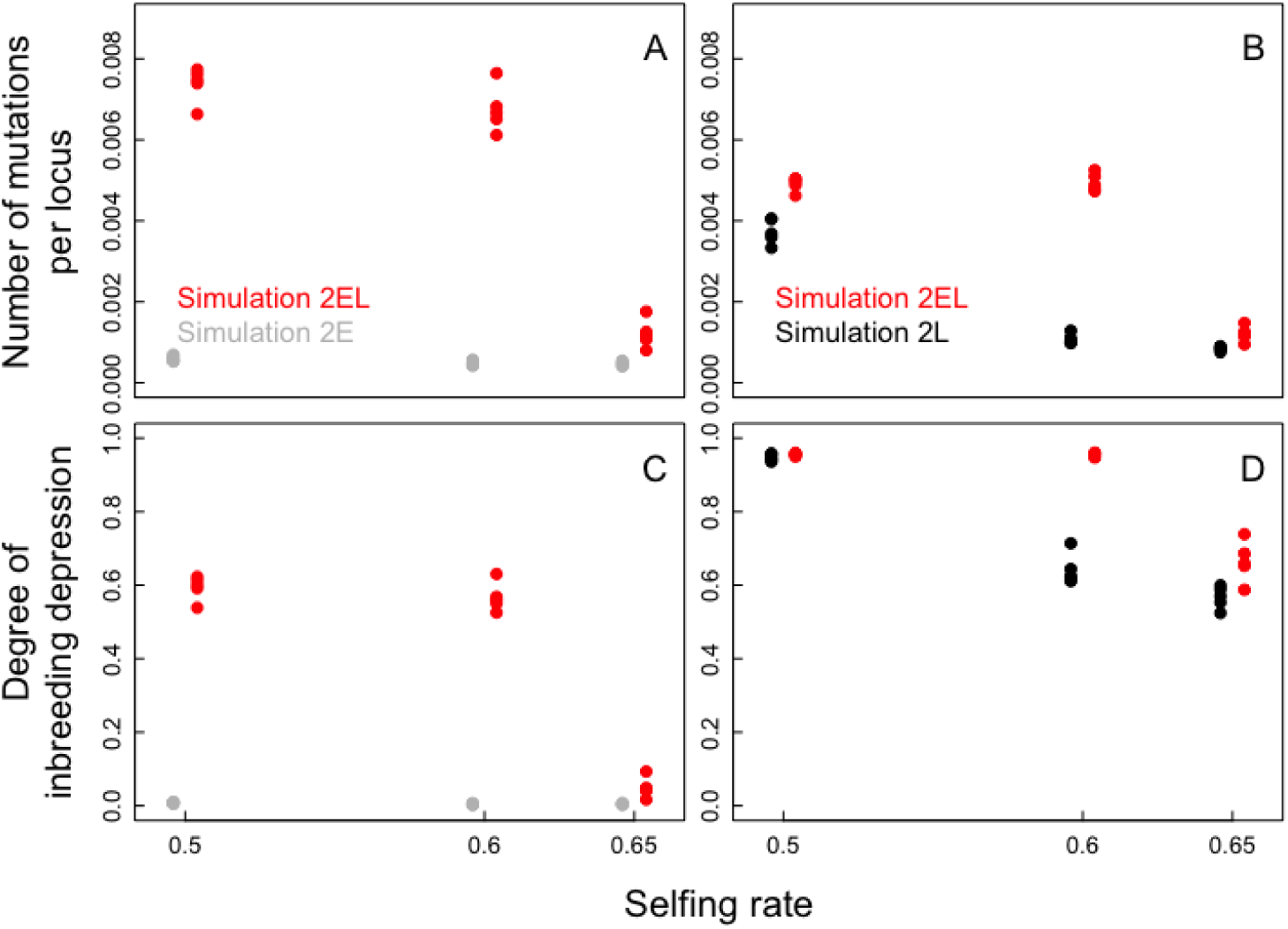
The results of the genetic architecture analysis in which each chromosome contains 200 early-acting loci and 1,800 late-acting loci (*n*_link_ = 5 and *n*_locus_ = 2000) (simulation 2). The dependences of the number of early- (A) and late-acting (B) mutations per locus after 500 generations and the degree of inbreeding depression related to the number of seeds produced (C) and the seed competitive ability (D) on the selfing rate, *s*, are shown. In all panels*, d*_e_ = 1 and *d*_l_ = 0.5 if early- and late-acting mutations occur, respectively. Red circles; simulation 2_EL_, in which mutations occur in both the early- and late-acting loci. Gray circles; simulation 2_E_, in which mutations occur in the early-acting loci but not in the late-acting loci. Black circles; simulation 2_L_, in which mutations occur in the late-acting loci but not in the early-acting loci. The simulation results of 5 runs are shown for each combination of parameter values.

**Figure S5.**
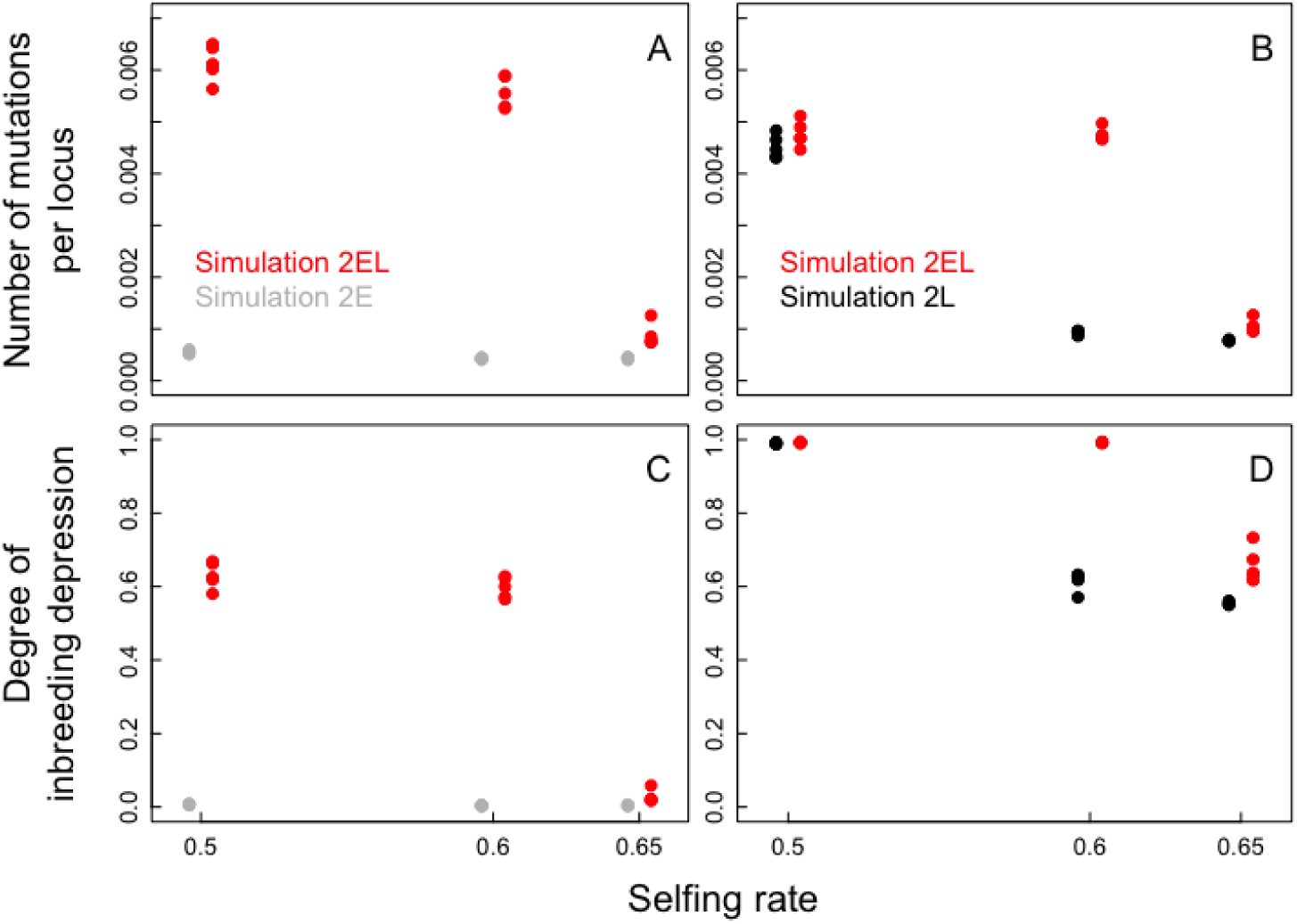
The results of the genetic architecture analysis in which all loci are unlinked, all genes segregate independently, and 1,000 early-acting loci and 9,000 late-acting loci are included (simulation 2). The dependences of the numbers of early-(A) and late-acting (B) mutations per locus after 500 generations and the degree of inbreeding depression related to the number of seeds produced (C) and the seed competitive ability (D) on the selfing rate, *s*, are shown. In all panels*, d*_e_ = 1 and *d*_l_ = 0.5 if early- and late-acting mutations occur, respectively. Red circles; simulation 2_EL_, in which mutations occur in both the early- and late-acting loci. Gray circles; simulation 2_E_, in which mutations occur in the early-acting loci but not in the late-acting loci. Black circles; simulation 2_L_, in which mutations occur in the late-acting loci but not in the early-acting loci. The simulation results of 5 runs are shown for each combination of parameter values.

**Figure S6.**
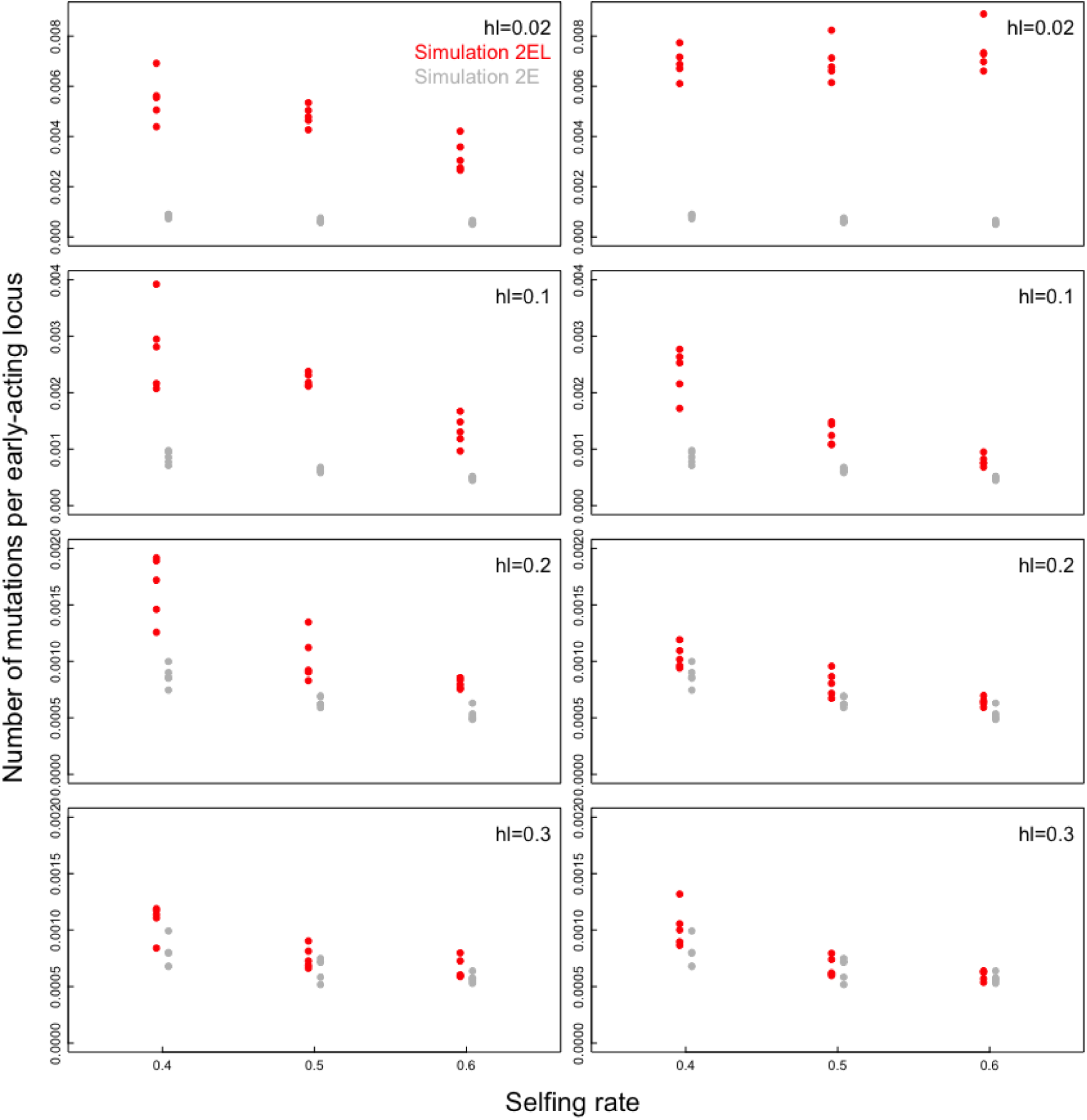
The effects of the dominance coefficient in the late-acting loci on the number of early-acting mutations maintained (simulation 2). The dependences of the number of early-acting mutations per locus after 500 generations on the selfing rate, *s*, the dominance coefficient in the late-acting loci, *h*_l_, and the selection coefficient against an individual late-acting mutation, *d*_l_ (*d*_l_ = 0.05 and 0.5 in the left and right panels, respectively), are shown. The early-acting mutations are lethal (*d*_e_ = 1), and *h*_e_ = 0.02 in all panels. Red circles; simulation 2_EL_, in which mutations occur in both the early- and late-acting loci. Gray circles; simulation 2_E_, in which mutations occur in the early-acting loci but not in the late-acting loci. The simulation results of 5 runs are shown for each combination of parameter values.

**Figure S7.**
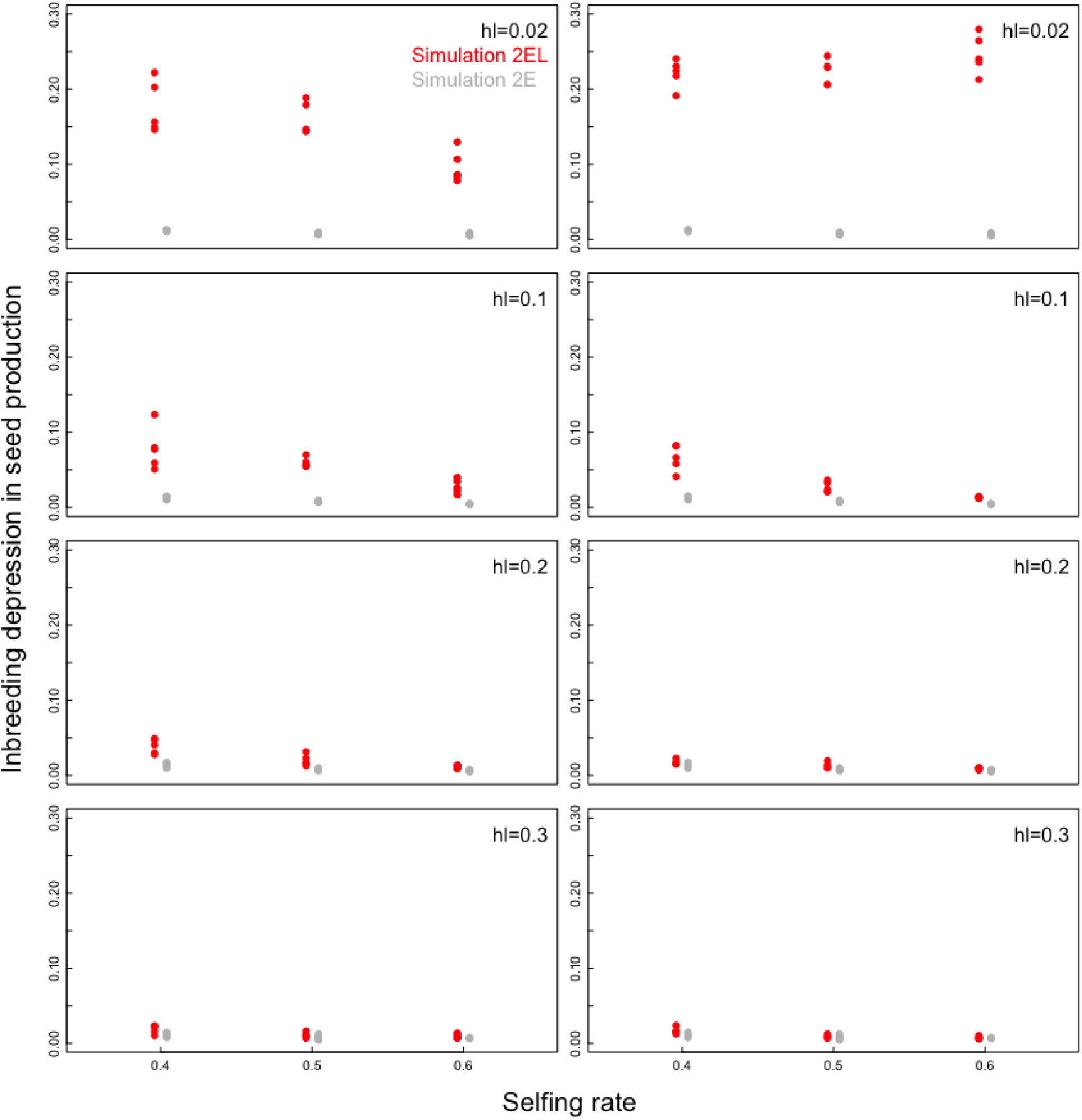
The effects of the dominance coefficient in the late-acting loci on the inbreeding depression in seed production (simulation 2). The dependences of the effects of inbreeding depression on seed production after 500 generations on the selfing rate, *s*, the dominance coefficient in the late-acting loci, *h*_l_, and the selection coefficient against an individual late-acting mutation, *d*_l_ (*d*_l_ = 0.05 and 0.5 in the left and right panels, respectively), are shown. The early-acting mutations are lethal (*d*_e_ = 1), and *h*_e_ = 0.02 in all panels. Red circles; simulation 2_EL_, in which mutations occur in both the early- and late-acting loci. Gray circles; simulation 2_E_, in which mutations occur in the early-acting loci but not in the late-acting loci. The simulation results of 5 runs are shown for each combination of parameter values.

**Figure S8.**
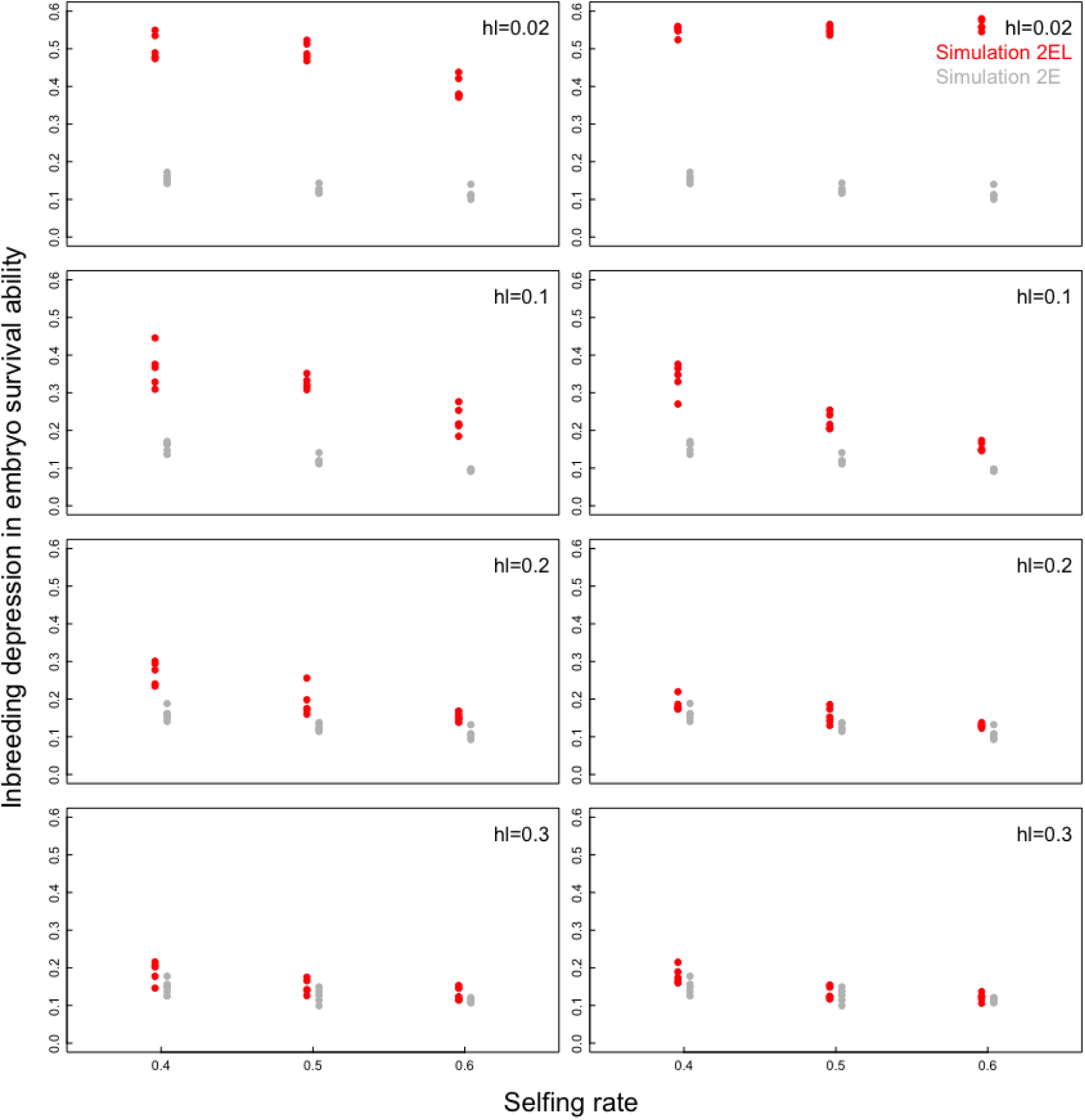
The effects of the dominance coefficient in the late-acting loci on the inbreeding depression in embryo survival ability (simulation 2). The dependences of the effects of inbreeding depression on the embryo survival ability after 500 generations on the selfing rate, *s*, the dominance coefficient in the late-acting loci, *h*_l_, and the selection coefficient against an individual late-acting mutation, *d*_l_ (*d*_l_ = 0.05 and 0.5 in the left and right panels, respectively), are shown. The early-acting mutations are lethal (*d*_e_ = 1), and *h*_e_ = 0.02 in all panels. Red circles; simulation 2_EL_, in which mutations occur in both the early- and late-acting loci. Gray circles; simulation 2_E_, in which mutations occur in the early-acting loci but not in the late-acting loci. The simulation results of 5 runs are shown for each combination of parameter values.

**Figure S9.**
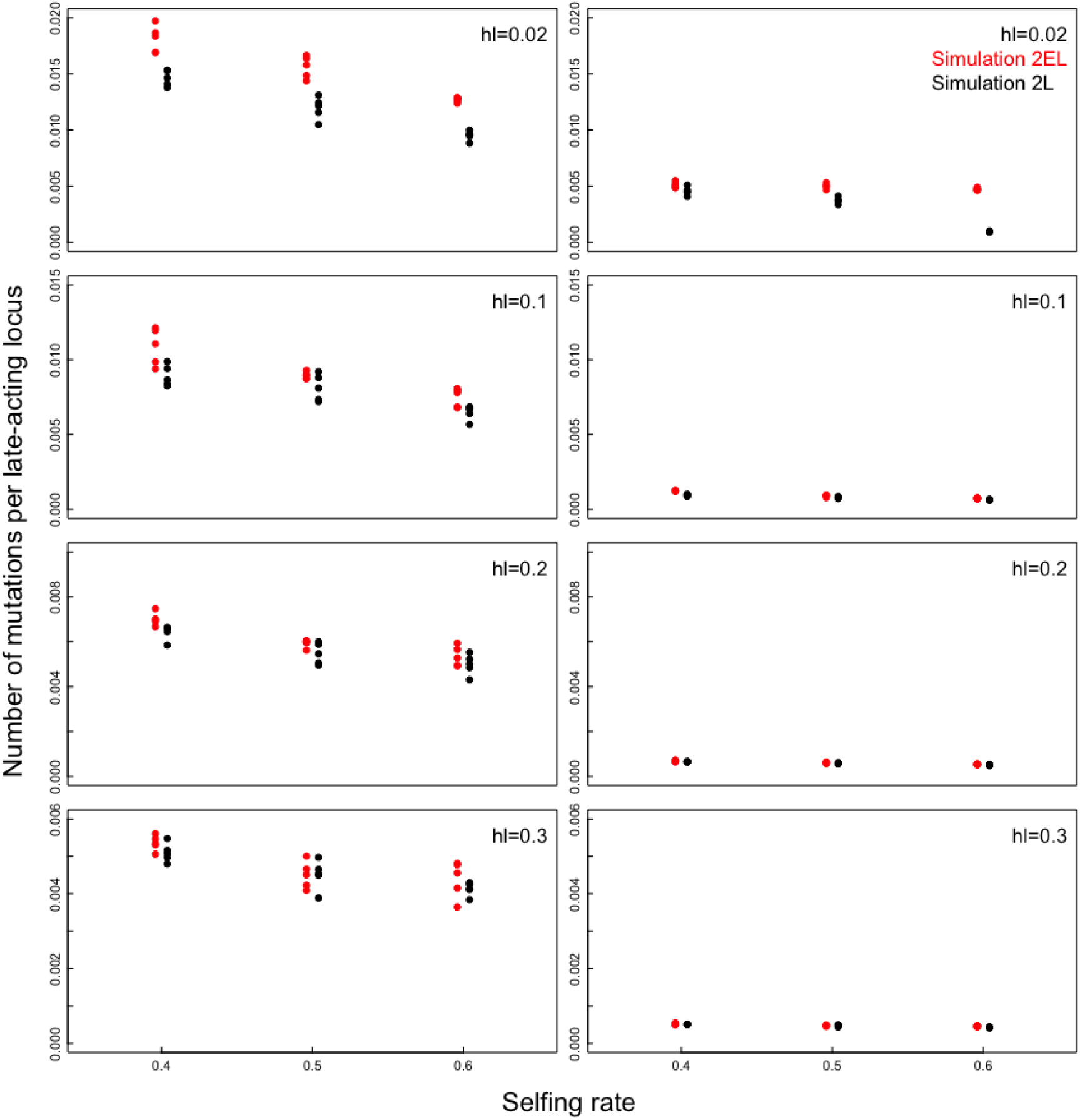
The effects of the dominance coefficient in the late-acting loci on the number of late-acting mutations maintained (simulation 2). The dependences of the number of late-acting mutations per locus after 500 generations on the selfing rate, *s*, the dominance coefficient in the late-acting loci, *h*_l_, and the selection coefficient against an individual late-acting mutation, *d*_l_ (*d*_l_ = 0.05 and 0.5 in the left and right panels, respectively), are shown. The early-acting mutations are lethal (*d*_e_ = 1), and *h*_e_ = 0.02 in all panels. Red circles; simulation 2_EL_, in which mutations occur in both the early- and late-acting loci. Black circles; simulation 2_L_, in which mutations occur in the late-acting loci but not in the early-acting loci. The simulation results of 5 runs are shown for each combination of parameter values.

**Figure S10.**
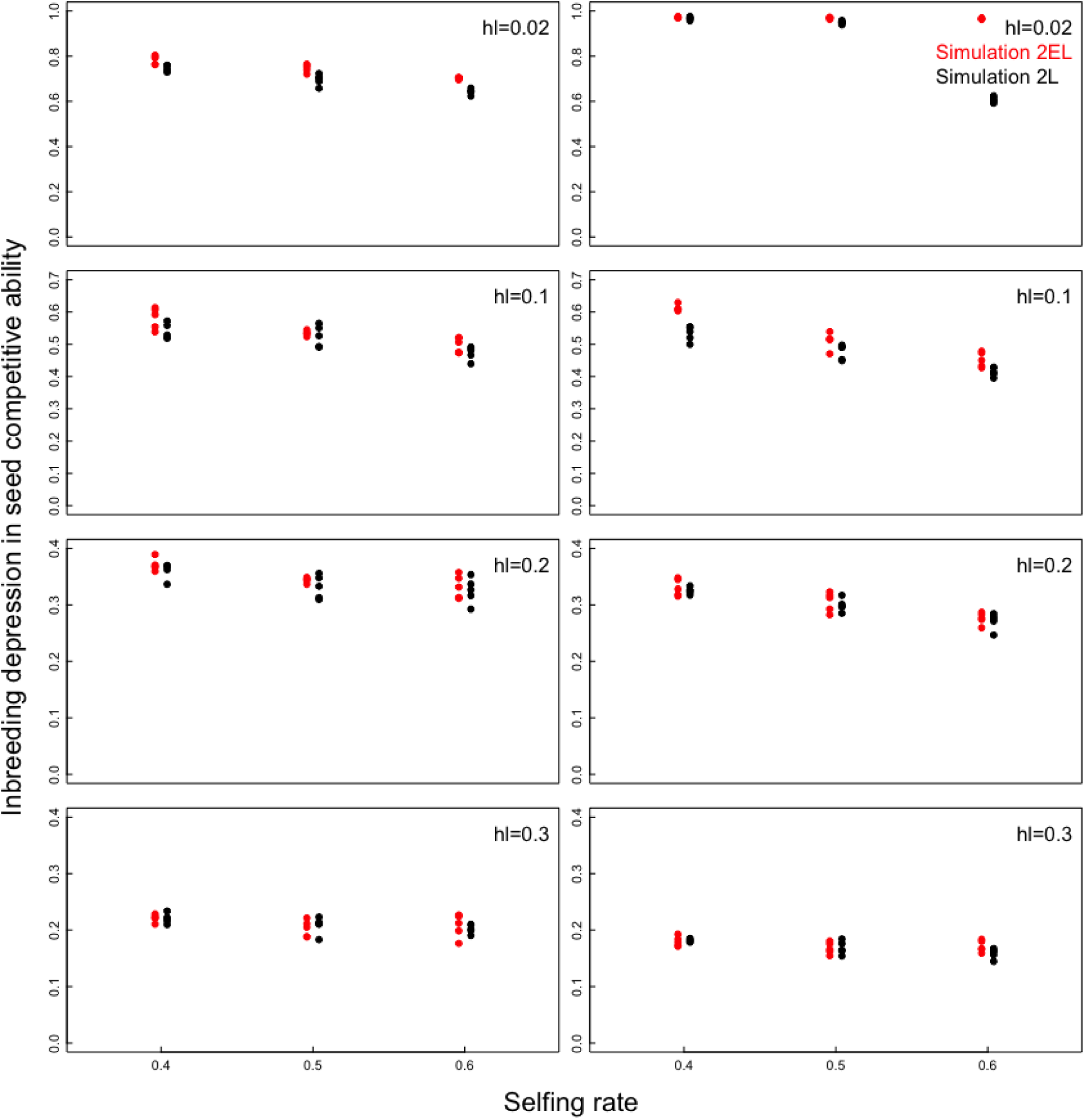
The effects of the dominance coefficient in the late-acting loci on the inbreeding depression in seed competitive ability (simulation 2). The dependences of the effects of inbreeding depression on the seed competitive ability after 500 generations on the selfing rate, *s*, the dominance coefficient in the late-acting loci, *h*_l_, and the selection coefficient against an individual late-acting mutation, *d*_l_ (*d*_l_ = 0.05 and 0.5 in the left and right panels, respectively), are shown. The early-acting mutations are lethal (*d*_e_ = 1), and *h*_e_ = 0.02 in all panels. Red circles; simulation 2_EL_, in which mutations occur in both the early- and late-acting loci. Black circles; simulation 2_L_, in which mutations occur in the late-acting loci but not in the early-acting loci. The simulation results of 5 runs are shown for each combination of parameter values.

## Literature Cited

Baldwin, S. J., and D. J. Schoen, 2019 Inbreeding depression is difficult to purge in self- incompatible populations of *Leavenworthia alabamica*. New Phytologist.

Barrett, S. C. H., 2002 The evolution of plant sexual diversity. Nature Reviews Genetics 3: 274–284.

Burd, M., 1998 “Excess” flower production and selective fruit abortion: a model of potential benefits. Ecology 79: 2123–2132.

Burd, M., and H. S. Callahan, 2000 What does the male function hypothesis claim? Journal of Evolutionary Biology 13: 735–742.

Charlesworth, B., and D. Charlesworth, 1999 The genetic basis of inbreeding depression. Genetics Research 74: 329–340.

Charlesworth, D., and J. H. Willis, 2009 The genetics of inbreeding depression. Nature Reviews Genetics 10: 783–796.

Cheptou, P. O., 2019 Does the evolution of self-fertilization rescue populations or increase the risk of extinction? Annals of Botany 123: 337–345.

Cheptou, P. O., and A. Mathias, 2001 Can varying inbreeding depression select for intermediary selfing rates? American Naturalist 157: 361–373.

Darwin, C., 1876 The Effects of Cross and Self Fertilisation in the Vegetable Kingdom. John Murray, London.

Eckert, C. G., K. E. Samis and S. Dart, 2006 Reproductive assurance and the evolution of uniparental reproduction in flowering plants, pp. 183–200 in Ecology and evolution of flowers, edited by L. D. Harder and S. C. H. Barrett. Oxford University Press, Oxford, UK.

Fisher, R. A., 1941 Average excess and average effect of a gene substitution. Annals of Eugenics 11: 53–63.

Fry, J. D., and S. V. Nuzhdin, 2003 Dominance of mutations affecting viability in Drosophila melanogaster. Genetics 163: 1357–1364.

Ganders, F. R., 1972 Heterozygosity for recessive lethals in homosporous fern populations: Thelypteris palustris and Onoclea sensibilis. Biological Journal of the Linnean Society 65: 211–221.

Goldberg, E. E., J. R. Kohn, R. Lande, K. A. Robertson, S. A. Smith et al., 2010 Species selection maintains self-incompatibility. Science 330: 493–495.

Goodwillie, C., S. Kalisz and C. G. Eckert, 2005 The evolutionary enigma of mixed mating systems in plants: occurrence, theoretical explanations, and empirical evidence. Annual Review of Ecology, Evolution, and Systematics 36: 47–79.

Holsinger, K. E., 1991 Mass-action models of plant mating systems: the evolutionary stability of mixed mating systems. American Naturalist 138: 606–622.

Huber, C. D., A. Durvasula, A. M. Hancock and K. E. Lohmueller, 2018 Gene expression drives the evolution of dominance. Nature Communications 9.

Kaul, S., H. L. Koo, J. Jenkins, M. Rizzo, T. Rooney et al., 2000 Analysis of the genome sequence of the flowering plant Arabidopsis thaliana. Nature 408: 796–815.

Kelly, J. K., 2007 Mutation–selection balance in mixed mating populations. Journal of theoretical Biology 246: 355–365.

Kozlowski, J., and S. C. Stearns, 1989 Hypotheses for the production of excess zygotes: models of bet-hedging and selective abortion. Evolution 43: 1369–1377.

Lande, R., and E. Porcher, 2015 Maintenance of quantitative genetic variance under partial self-fertilization, with implications for evolution of selfing. Genetics 200: 891–906.

Lande, R., and E. Porcher, 2017 Inbreeding depression maintained by recessive lethal mutations interacting with stabilizing selection on quantitative characters in a partially self-fertilizing population. Evolution 71: 1191–1204.

Lande, R., and D. W. Schemske, 1985 The evolution of self-fertilization and inbreeding depression in plants. I. genetic models. Evolution 39: 24–40.

Lande, R., D. W. Schemske and S. T. Schultz, 1994 High inbreeding depression, selective interference among loci, and the threshold selfing rate for purging recessive lethal mutations. Evolution 48: 965–978.

Latta, R., and K. Ritland, 1994 Conditions favoring stable mixed mating systems with jointly evolving inbreeding depression. Journal of theoretical Biology 170: 15–23.

Lee, T. D., 1988 Patterns of fruit and seed production, pp. 179–202 in Plant reproductive ecology, edited by J. Lovett-Doust and L. Lovett-Doust. Oxford University Press, New York.

Lloyd, D. G., 1979 Some reproductive factors affecting the selection of self fertilization in plants. American Naturalist 113: 67–79.

Lloyd, D. G., 1992 Self- and cross-fertilization in plants. II. the selection of self-fertilization. International Journal of Plant Sciences 153: 370–380.

Lloyd, D. G., and D. J. Schoen, 1992 Self- and cross-fertilization in plants. I. Functional dimensions. International Journal of Plant Sciences 153: 358–369.

Peters, A. D., D. L. Halligan, M. C. Whitlock and P. D. Keightley, 2003 Dominance and overdominance of mildly deleterious induced mutations for fitness traits in *Caenorhabditis elegans*. Genetics 165: 589–599.

Porcher, E., and R. Lande, 2005a The evolution of self-fertilization and inbreeding depression under pollen discounting and pollen limitation. Journal of Evolutionary Biology 18: 497–508.

Porcher, E., and R. Lande, 2005b Reproductive compensation in the evolution of plant mating systems. New Phytologist 166: 673–684.

Roze, D., 2015 Effects of interference between selected loci on the mutation load, inbreeding depression, and heterosis. Genetics 201: 745–757.

Sato, S., S. Tabata, H. Hirakawa, E. Asamizu, K. Shirasawa et al., 2012 The tomato genome sequence provides insights into fleshy fruit evolution. Nature 485: 635–641.

Schemske, D. W., and R. Lande, 1985 The evolution of self-fertilization and inbreeding depression in plants. II. Empirical observations. Evolution 39: 41–52.

Stebbins, G. L., 1974 Flowering plants: evolution above the species level. Harvard University Press, Cambridge.

Sutherland, S., 1987 Why hermaphroditic plants produce many more flowers than fruits: experimental tests with *Agave mckelveyana*. Evolution 41: 750–759.

Szafraniec, K., D. M. Wloch, P. Sliwa, R. H. Borts and R. Korona, 2003 Small fitness effects and weak genetic interactions between deleterious mutations in heterozygous loci of the yeast Saccharomyces cerevisiae. Genetical Research 82: 19–31.

Uyenoyama, M. K., and D. M. Waller, 1991 Coevolution of self-fertilization and inbreeding depression. I. Mutation-selection balance at one and two loci. Theoretical Population Biology 40: 14–46.

Winn, A. A., E. Elle, S. Kalisz, P.-O. Cheptou, C. G. Eckert et al., 2011 Analysis of inbreeding depression in mixed-mating plants provides evidence for selective interference and stable mixed mating. Evolution 65: 3339–3359.

